# Skeletal Muscle Stem Cells Modulate Niche Function in Duchenne Muscular Dystrophy through YY1-CCL5 Axis

**DOI:** 10.1101/2024.01.13.575317

**Authors:** Yang Li, Chuhan Li, Qiang Sun, Fengyuan Chen, Yeelo Cheung, Yu Zhao, Ting Xie, Bénédicte Chazaud, Hao Sun, Huating Wang

**Author notes:** Co-correspondence: Huating Wang, 507A Li Ka Shing Institute of Health Sciences, Prince of Wales Hospital, Chinese University of Hong Kong, Hong Kong SAR, China.

## Abstract

Stem cell activity is known to be tightly regulated by both intrinsic and extrinsic pathways but less is known about whether and how stem cells modulate their niche microenvironment. Adult skeletal muscle stem cells (MuSCs) are indispensable for muscle regeneration and also tightly regulated by macrophages (MPs) and fibro-adipogenic progenitors (FAPs) in the niche. Deregulated MuSC/MP/FAP interactions and the ensuing inflammation and fibrosis are hallmarks of dystrophic muscle. Here in this study we demonstrate that intrinsic deletion of transcription factor YY1 in MuSCs exacerbates dystrophic pathologies by altering the cellular composition and heterogeneity of MPs and FAPs. Further analysis reveals that the YY1 loss induces the expression of immune genes in MuSCs, including *Ccl5*. Augmented secretion of CCL5 from MuSCs promotes the recruitment of MPs via CCL5/CCR5 mediated crosstalk, which subsequently hinders the apoptosis and clearance of FAPs through elevated TGFβ1 accumulation. Maraviroc mediated pharmacological blockade of the CCL5/CCR5 axis effectively mitigates muscle dystrophy and improves muscle performance. Lastly, we further demonstrate that YY1 represses *Ccl5* transcription in MuSCs by directly binding to its enhancer thus facilitating promoter-enhancer looping. Altogether, our study has demonstrated the previously unappreciated role of MuSCs in actively shaping their niche microenvironment through secreting immunomodulatory cytokines, and has also provided novel insight into the therapeutic intervention of muscle dystrophy.

## Introduction

Skeletal muscle has a robust regenerative capacity, with rapid re-establishment of full power occurring even after severe damage that causes widespread myofiber necrosis. The cells responsible for muscle regeneration are adult muscle stem cells (MuSCs, also called satellite cells) which are located in a niche beneath the ensheathing basal lamina on the surface of the myofibers in a quiescent stage under normal conditions(*1–3*). Upon injury, MuSCs are rapidly activated to become myoblasts, undergo proliferative expansion, and eventually differentiate into myotubes and fuse to form new myofibers(*1–3*). A subset of MuSCs undergoes self-renewal and returns to the quiescent state to replenish the adult stem cell pool. Each phase of the activities is tightly orchestrated at two levels. First, through intrinsic preprogrammed mechanisms and, second, through extrinsic regulations imposed by the stem cell microenvironment or so-called stem cell niche. The intrinsic regulatory mechanisms have been relatively well defined. For example, it is widely accepted that gene regulation at transcriptional level by transcription factors (TFs) plays crucial roles in instructing MuSC regenerative responses. Among many key TFs, Yin Yang 1(YY1) is ubiquitously expressed but possesses unique transcriptional regulatory functions in MuSCs based on findings from our group and others(*4–9*). For example, we recently elucidated the function of YY1 in acute injury-induced muscle regeneration as a key regulator of MuSC activation/proliferation through its dual roles in modulating metabolic pathways(*4*).

MuSC functionality is also tightly controlled by the crosstalk between MuSCs and other cell types within their niche(*10*). The MuSC niche is relatively static under homeostatic conditions to maintain MuSC quiescence and undergo dynamic remodeling following injury through a spatiotemporally tightly coordinated flux of different cell types such as inflammatory, vascular, and mesenchymal cells. The reciprocal functional interactions among these cell types are crucial in coordinating the repair of injured muscles. In particular, optimal regeneration entails a sequence of events that ensure temporally coordinated interactions among MuSCs, macrophages (MPs) and fibro-adipogenic progenitors (FAPs)(*11–13*). An initial recruitment of MPs is typically followed by the sequential activation of FAPs and MuSCs. The key immune and non-immune functions MPs exert in muscle regeneration are well accepted(*14, 15*). The pro-inflammatory response following acute muscle injury usually begins with infiltration of neutrophils, followed by pro-inflammatory MPs featured by Ly6C^high^F4/80^low^, which produces mostly inflammatory cytokines, such as TNFα, IL-1β, and IFNγ to promote MuSC activation and proliferation. Afterward, at later stages of regeneration, a distinct Ly6C^low^F4/80^high^ pro-regenerative and anti-inflammatory MPs become more prevalent, which produce different cytokines, such as IL-4, IL-10 and TGFβ1, to promote myoblast differentiation and tissue repair(*14, 16*). Although much is known about the impact of MPs on MuSCs, the possible reciprocal regulation of MuSCs on MPs remains largely unexplored despite a pioneer work long time ago demonstrating that human myoblasts can indeed secrete an array of chemotactic factors to initiate monocyte chemotaxis in culture(*17*). FAPs are a muscle interstitial mesenchymal cell population which can differentiate into fibroblasts, adipocytes and possibly into osteoblasts and chondrocytes. Emerging evidence solidifies FAPs’ critical role in efficient muscle repair(*12*). Upon muscle injury, FAPs become activated, proliferate and expand, and provide a transient favorable microenvironment to promote MuSC-mediated regeneration. FAP expansion is critical during regeneration to sustain MuSC proliferation and differentiation in a paracrine manner and maintain the MuSC pool, however, as regeneration proceeds, their timely removal from the regenerative niche through apoptosis is necessary to prevent pathological accumulation and muscle dysfunction(*18*). Again, it remains largely unknown if FAPs receive any reciprocal signaling from MuSCs. Nevertheless, it is widely accepted that finely tuned molecular interactions of FAPs and MPs are essential for successful regeneration. Expansion and decline of FAP cell populations following injury are determined by MPs and disrupted MP dynamics can result in aberrant retention of FAPs in muscles following acute or chronic injuries(*11, 12*). For example (*18*) in acutely damaged skeletal muscle, infiltrating MPs, through their expression of TNFα, directly induce apoptosis and the timely clearance of FAPs, thus preventing the occurrence of fibrosis.

Therefore, the finely orchestrated functional interactions among MuSCs, MPs and FAPs are crucial to instruct the proper progress of damage-induced muscle repair. Conditions that compromise the functional integrity of this network can skew muscle repair toward pathological outcomes, for example in the case of Duchene muscular dystrophy (DMD)(*11, 13*), which is a lethal progressive pediatric muscle disorder caused by the genetic mutations in the dystrophin gene. In the absence of dystrophin protein, DGC (dystrophin-glycoprotein complex) assembly on the muscle member is impaired which weakens the muscle fibers, rending them highly susceptible to injury. Muscle contraction-induced stress results in constant cycles of degeneration and regeneration, resulting in chronic inflammation and fibro-fatty tissue replacement. Recent advances in single cell RNA-sequencing (scRNA-seq) have enabled us to characterize cell compositions and interactions in both human and mouse DMD muscles(*19, 20*). Compared to muscles underlying acute injury-regeneration, more intricate cellular dynamics and interactions are observed by dystrophic muscles. An increased prevalence of MPs and FAPs was confirmed and correlated with disease severity. MPs are highly heterogeneous and partially pathogenic in dystrophic muscle(*21*). A plethora of MP-derived factors are critically involved in inflammation and fibrosis of the muscle. It is speculated that the heterogeneous and pathogenic activation of MPs in dystrophic muscle is likely induced by the asynchronous regeneration altered microenvironment but the underlying signaling molecules and cellular sources remain unexplored(*22*). Evidence from the scRNA-seq profiling also revealed remarkable heterogeneity of FAPs and their correlation with disease severity(*19*). FAPs undergo uncontrolled expansion and resistance to clearance in dystrophic muscle, which plays a prominent role in intramuscular fat deposition and fibrosis(*23*). Increased TGFβ signaling is believed to prevent FAP apoptosis and induces their differentiation into matrix-producing cells to cause fibrosis in dystrophic muscle(*18*).

Overall, cellular communications among the main pathogenic cells in dystrophic muscle warrant further investigation; in particular, it is unknown if MuSCs play an active role in initiating cross talks with MPs and FAPs to manipulate their own niche microenvironment. In fact, it is safe to state that in general our understanding about if and how tissue adult stem cells impact their niches is very limited. Across a variety of well-studied adult stem cells such as mesenchymal stem cells (MSCs), hematopoietic stem cells (HSs) and neural stem cells (NSCs) etc., we have learnt plenty about the essential roles of the stem cell niche in regulating stem cell behavior and functionality(*24*), including how alteration in the stem cell niche causes cellular damage and impairs the regenerative capacity of stem cells. In principle the stem cells are perceived as the passive recipient of niche signals and impact. It is rare to find studies on if and how stem cells can actively contribute to the niche integrity in homeostasis and how the intrinsic changes in stem cells are connected to extrinsic niche alterations in pathological conditions. Here in this study, we discovered that intrinsic deletion of YY1 in MuSCs of dystrophic mdx mice (dKO) exacerbated fibrosis and inflammation. Analysis of cellular compositions uncovered elevated numbers of MPs and FAPs accompanying a decrease in MuSCs in dKO vs. control mice; moreover, scRNA-seq profiling revealed altered cellular heterogeneity of MPs and FAPs. Furthermore, we found that YY1 deletion in MuSCs induced up-regulation of immune genes thus rendering MuSCs immunogenic. Notably, CCL5 was identified as a critical factor in facilitating the recruitment of MPs via the CCL5/CCR5 axis mediated MuSC-MP interaction; Escalated MP accumulation subsequently prevented FAP apoptosis and clearance via increased TGFβ accumulation. Consistently, treatment of the dKO mice with a CCR5 antagonist Maraviroc (MVC) significantly ameliorated the dystrophic pathologies and muscle function in dKO mice. Lastly, we elucidated that *Ccl5* induction in dKO MuSCs was resulted froman altered enhancer-promoter loop interaction. Altogether our findings demonstrate an active role of MuSCs in orchestrating cellular interactions with other niche cells and highlight their previously unappreciated capacity in modulating niche microenvironment through their immune-secretory function.

## Results

### Inducible deletion of YY1 from MuSCs aggravates muscle dystrophy in mdx mouse

Our prior study(*4*) has demonstrated that YY1 plays an indispensable role in acute injury-induced muscle regeneration and also hinted its involvement in chronic degeneration/regeneration occurring in dystrophic mdx mice. To further elucidate the role of YY1 in dystrophic muscle, we characterized the phenotypes of the YY1/mdx double knockout (dKO) mice that were generated by inducible deletion of YY1 in MuSCs of mdx mice (Suppl. Fig. S1A). Five days of consecutive intraperitoneal (IP) administration of tamoxifen (TMX) was performed in 1.5-month (1.5M)-old Ctrl and dKO mice to induce YY1 deletion in MuSCs and the muscles were harvested when mice were 2M, 2.5M, 3.5M and 8.5M old for subsequent analyses (Fig. 1A-D). Consistent with our prior observations(*4*), YY1 deletion caused severe impairment in muscle regeneration in both limb (tibialis anterior, TA) and diaphragm (DP) muscles as evidenced by the abnormal hypertrophy of muscle fibers assessed by H&E (Fig. 1E) and eMyHC (Suppl. Fig. S1B-C) staining at 2.5M. And a decreased number of MuSCs (Suppl. Fig. S1D) caused by impaired proliferation (measured by in vivo EdU incorporation assay in Suppl. Fig. S1E) was also confirmed. Moreover, the dKO mice displayed striking fibrotic-and immune-phenotypes. An excessive amount of fibrosis was detected in dKO by both Masson’s trichrome staining (Fig. 1F) and increased expression of a panel of fibrotic markers (Suppl. Fig. S1F-G). The level of fibrosis appeared to arise from increased number of PDGFRα + FAPs as a positive correlation was detected between the main fluorescence intensity (MFI) of PDGFRα and COL1a1 protein staining in both TA (Fig. 1G) and DP (Fig. 1H) of Ctrl and dKO mice. This was accompanied by an elevated level of inflammation as indicated by increased expression of inflammatory markers (Suppl. Fig. S1H-I) and an augmented number of macrophages (Fig. 1I-L). Altogether, the above findings demonstrate that YY1 deletion in MuSCs leads to exacerbation of dystrophic pathologies in mdx mice. The phenotypes persisted in dKO mice into elder age by contrasting with the Ctrl mdx mice alleviated at around 3.5 months-old (Suppl. Fig. S1J-M). As a result, the dKO mice were overall very fragile displaying much smaller body size and lower body weight (Fig. 1M-N) and muscle weight at 8.5 months; the TA muscle mass decreased by 50.97% (Fig. 1O) and the thickness of DP muscle shrunk by 39.51% (Fig. 1P). When subject to treadmill (Fig. 1Q) or voluntary wheel-running (Fig. 1R) exercises, these dKO mice exhibit poor performance compared to the Ctrl mice. As expected, the survival rate of the dKO mice was significantly reduced and half of them died before six months (Fig. 1S). Therefore, the YY1 loss in MuSCs worsens the dystrophic manifestations in the *mdx* mouse.

**Fig. 1.**
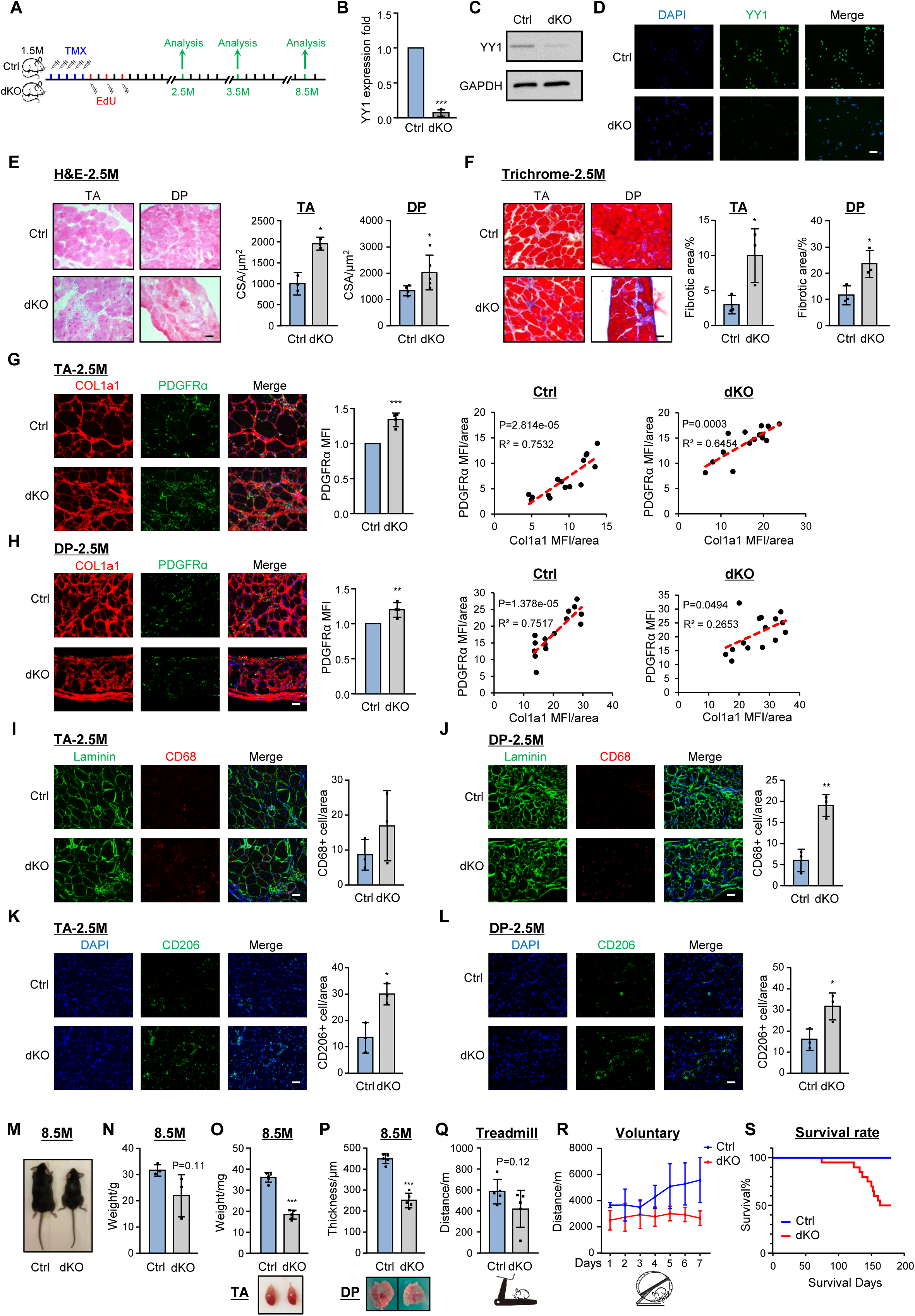
Inducible deletion of YY1 in MuSCs aggravates muscle dystrophy in mdx mouse. **A** Schematic of the experimental design for analyzing dystrophic phenotypes in Ctrl and dKO mice. **B-D** Validation of YY1 ablation in dKO MuSCs by RT–qPCR, Western blot and IF staining, n=3. **E** Left panel: H&E staining of TA and DP muscles collected from the Ctrl and dKO mice at the age of 2.5M. Scale bar: 50 μm. Right panel: quantification of cross-sectional areas (CSAs) of the stained fibers, n≥3. **F** Left panel: Masson’s Trichrome staining of TA and DP muscles collected from the Ctrl and dKO mice at the age of 2.5M. Scale bar: 50 μm. Right panel: quantification of the stained fibrotic areas, n=3. **G-H** IF staining of Collagen1a1 (COL1a1, red) and PDGFRα (green) was performed on the above TA and DP muscles from the Ctrl and dKO mice. Mean fluorescence intensity (MFI) of the stained sections was quantified, correlation of COL1a1 and PDGFRα MFI was calculated. Scale bar: 50 μm, n=4. **I-J** Left: IF staining of DAPI (blue), Laminin (green) and CD68 (red) on the above TA and DP muscles. Right: quantification of the number of CD68+ cells per area. Scale bar: 50 μm, n=3. **K-L** Left: IF staining of DAPI (blue) and CD206 (green) on the above TA and DP muscles. Right: quantification of the number of CD206+ cells per area. Scale bar: 50 μm, n=3. **M** Representative images of Ctrl and dKO mice at the age of 8.5M. **N-P** Body weight, TA muscle weight and DP muscle thickness were measured in the aforementioned mice, n≥3. **Q** 2.5M Ctrl and dKO mice were subject to treadmill exercise and the running distance until exhaustion is shown, n=5. **R** The above mice were subject to voluntary wheel-running exercise and the weekly running distance is shown, n=4. **S** Survival rate of Ctrl and dKO mice at 6 months), n=10. All the bar graphs are presented as mean ±SD, Student’s t test (two-tailed unpaired) was used to calculate the statistical significance (B, E–L, N-Q): *p < 0.05, **p < 0.01, ***p < 0.001, n.s. = no significance.

### Intrinsic YY1 deletion from MuSCs alters their niche in dystrophic muscle

We hypothesized that YY1 deletion in MuSCs exacerbates dystrophic phenotypes possibly by increasing the numbers of MPs and FAPs in the niche microenvironment while shrinking the MuSC pool. To test this hypothesis, MuSCs, FAPs and MPs were isolated from limb muscles at 5, 21, and 60 days after TMX administration (Fig. 2A) following established FACS sorting protocols(*4, 25, 26*) (Suppl. Fig. S2A-C). As expected, we observed a progressive decline of MuSC in the dKO but not in the Ctrl (Fig. 2B). Meanwhile, FAPs (Fig. 2C) and MPs (Fig. 2D) were significantly increased at both 21 and 60 days post TMX injection. The above results suggest that the intrinsic deletion of YY1 in MuSCs leads to skewed cellular composition in mdx muscle.

**Fig. 2.**
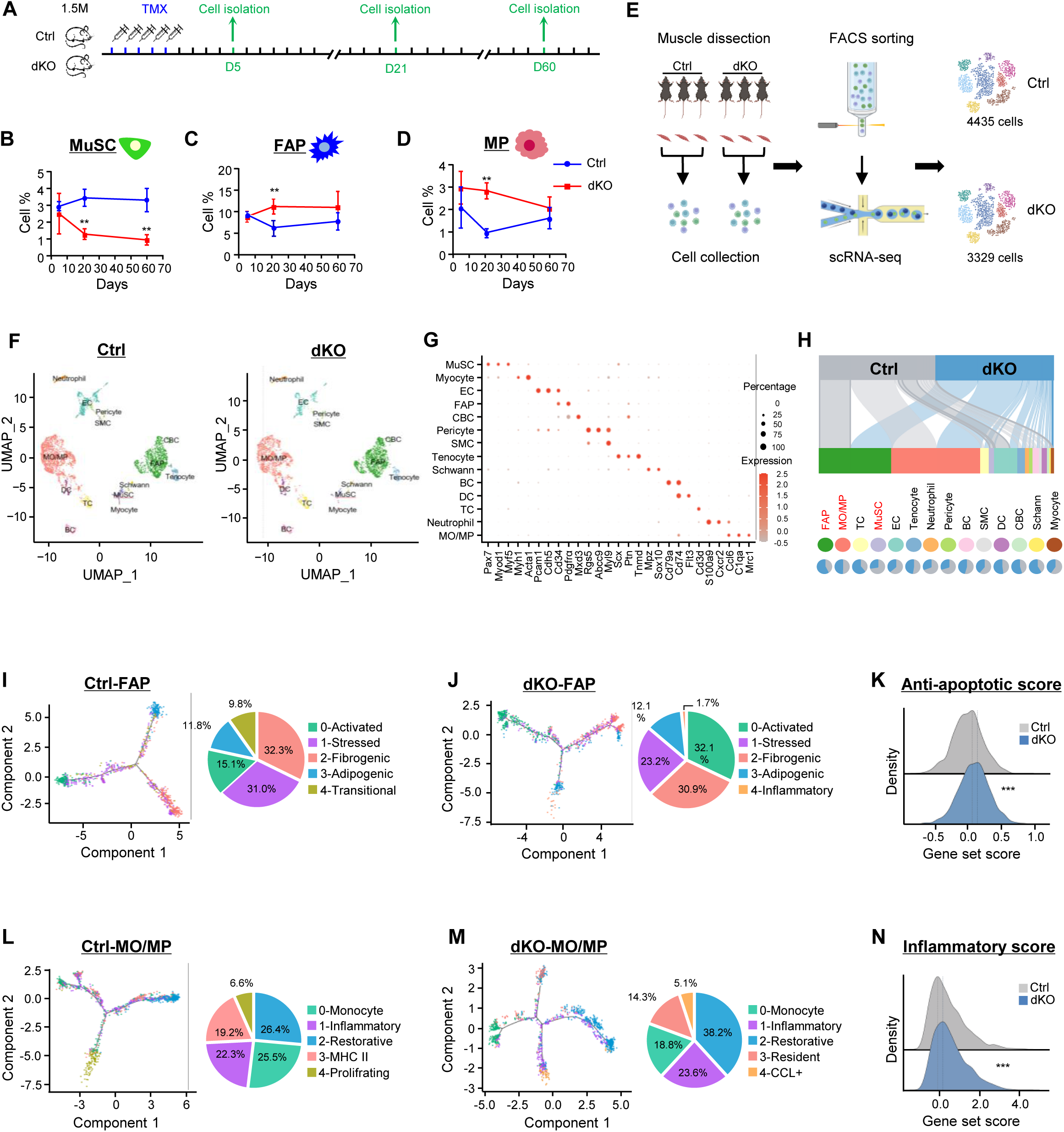
Intrinsic YY1 deletion in MuSCs alters cellular microenvironment in dystrophic muscle. **A** Schematic of the experimental design for analyzing MuSC, FAP and MP populations in Ctrl and dKO mice. **B-D** The percentages of isolated MuSCs, FAPs and MPs from Ctrl and dKO mice at 5, 21 and 60 days after TMX administration, n=3. **E** Schematic of scRNA-seq experimental design, mononuclear cells from three pairs of Ctrl and dKO mice were combined for FACS sorting and living cell selection. A total of 4435 and 3329 living cells from Ctrl and dKO mice were identified respectively. **F** scRNA-seq was performed in the whole muscle from three pairs of Ctrl and dKO mice. Data are shown as a uniform manifold approximation and projection (UMAP) to visualize variation in single-cell transcriptomes. Unsupervised clustering resolved at least 14 cell types (colour coded). **G** Dot plot showing the expression signatures of representative marker genes for each cell type. **H** Top: Sankey plots showing the distribution of Ctrl and dKO cells across different cell types. Bottom: pie plots showing the relative cell proportion between Ctrl and dKO groups across different cell types. **I-J** Left: pseudotime trajectory inference of the identified FAP subpopulations in Ctrl and dKO. Right: pie charts showing the relative cell proportion of each subtype. **K** Ridge map showing the global distribution density of anti-apoptotic of Ctrl and dKO FAPs. The corresponding dashed line represents the peak position of each group. **L-M** Left: pseudotime trajectory inference of the identified MO/MP subpopulations in Ctrl and dKO. Right: pie charts showing the relative cell proportion of each subtype. **N** Ridge map showing the global distribution density of inflammatory score of Ctrl and dKO MO/MP. The corresponding dashed line represents the peak position of each group. All the bar graphs are presented as mean ± SD, Student’s t test (two-tailed unpaired) was used to calculate the statistical significance (B-D, K, N): *p < 0.05, **p < 0.01, ***p < 0.001, n.s. = no significance.

To further illuminate the altered muscle niche we performed single cell (sc) RNA-seq. As shown in Fig. 2E, hind limb muscles were collected from three pairs of Ctrl or dKO mice (one month after TMX injection); a mixed single-cell suspension from the three mice were subject to droplet-based scRNA-seq on a 10x Chromium platform; 4335 and 3329 cells were obtained from Ctrl and dKO groups for subsequent analyses. A comprehensive definition of cellular atlas uncovered a total of 14 different cell populations including Monocyte/Macrophage (MO/MP), FAP, endothelial cell (EC), tenocyte, T cell (TC), B cell (BC), smooth muscle cell (SMC), dendritic cell (DC), MuSC, neutrophil, cycling basal cell (CBC), myocyte, pericyte and schwann cell (Fig. 2F) based on normalized gene expression levels and canonical cell type-specific markers (Fig. 2G, Suppl. Fig. S2J, and Suppl. Table S2). Notably, FAP, MO/MP were the major cell populations in the niche, making up over 60% of the total cells in both Ctrl and dKO (Fig. 2H). Consistent with the results from Fig. 2B-C, a remarkably increased population of FAPs was detected in dKO vs. Ctrl (35.9% vs. 26.4%) accompanied by a reduced population of MuSCs (1.2% vs. 3.3%). Interestingly, no obvious increase in the number of MO/MP was observed (38.0 % vs. 38.2%). The ratio of most cell populations, including EC (7.2% vs. 12.6%), neutrophil (1.1% vs. 2.6%) and pericyte (0.9% vs. 2.1%), displayed a decrease in dKO vs. Ctrl (Fig. 2H, Suppl. Table S2).

To further illuminate the dynamic shifts of FAPs, pseudotime trajectory was utilized to reveal the cellular heterogeneity and fate determination. Unbiased SNN clustering(*27*) uncovered five subpopulations of FAPs in Ctrl group and designated as 0-Activated, 1-Stressed, 2-Fibrogenic, 3-Adipogenic and 4-Transitional according to differentially expressed genes (DEGs) (Fig. 2I, Suppl. Fig. S2K, Suppl. Table S2). Activated FAPs localized toward the starting point of the trajectory, which were marked by the expression of *Cxcl5*, *Cxcl3*, *Ccl7*, and *Ccl2*; Stressed FAPs did not appear to have any spatial bias, and were enriched with stress-responsive genes such as heat shock genes (Hspb1, Hspd1, Hspe1) and AP-1 family transcription factors (*Fos*, *Atf3*, *Jun*); Fibrogenic (enriched with ECM factors, *Cxcl14*, *Lum*, *Smoc2*, *Lpl*) and Adipogenic FAPs (enriched with adipogenic factors, *Pi16*, *Dpp4*, *Fn1)* (Supp. Fig. S2K) highlighted the main divergent fates of two FAP subpopulations in the trajectory. Transitional FAPs appeared on the path from the Activated to Fibrogenic/Adipogenic FAPs, marked with *Apod*, *Ptx3*, *Myoc* and *Mt1* genes. As comparison, above-described five subsets of FAPs were also identified in dKO muscles including 0-Activated, 1-Stressed, 2-Fibrogenic, 3-Adipogenic and 4-Inflammatory (Fig. 2J, Suppl. Fig. S2L, and Suppl. Table S2). And a similar trajectory was adopted: starting from the Activated state, dKO FAPs were arranged along a trajectory that diverged into two distinct branches, which coincided with the two subpopulations, Fibrogenic and Adipogenic (Fig. 2J). Notably, the proportion of the Activated subset increased significantly in dKO vs. Ctrl (32.1% vs. 15.1%), while the Stressed subset displayed a decrease (31.0% vs. 23.2%) (Fig. 2I-J). According to a previous study(*28*), activated FAPs emerge in the early stage of muscle injury, therefore the phenomenon was in line with the delayed regeneration in dKO. Meanwhile, stressed FAPs were enriched with heat shock genes, which were closely related with cellular apoptosis(*29, 30*), suggesting resistance of apoptosis in the dKO FAPs. Indeed, a significantly higher anti-apoptotic score (defined by anti-apoptotic gene set, Supp. Table S2) was measured in dKO vs. Ctrl FAPs (0.16 vs. 0.08) (Fig. 2K) alongside a decreased apoptotic score (Suppl. Fig. S2M, Suppl. Table S2). The above results suggest that enhanced apoptosis resistance may explain the elevated FAP population in dKO muscle niche. It is also interesting to point out that an inflammatory FAP subset, marked by elevated expression of phagosome and chemokine pathway related genes, was identified exclusively in the dKO but not in Ctrl group (Fig. 2I-J, Suppl. Fig. S2K-L).

When examining MO/MP subpopulations and heterogeneity, we identified five subsets in Ctrl muscle defined as 0-Monocyte, 1-Inflammatory, 2-Restorative, 3-MHC II, 4-Proliferating (Fig. 2L, Suppl. Table S2)(*28, 31*). Monocytes were characterized by highly expressed *Cxcl3*, *Vcan* and *Chil3*; Inflammatory MPs were enriched for *Spp1*, *Fabp5* and *Cd36*; Restorative MPs expressed high levels of *C1qa*, *C1qb*, and *C1qc*; MHC II MPs were harbored an abundance of *H2-Aa*, *H2-Eb1*, and *H2-Ab1*; The Proliferating subset was distinguished by highly expressed cell cycle and cell division related genes such as *Ccna2*, *Ccnb2*, *Cdk1*, *Cdc20*, *Cdca3*, *Cdca8* (Suppl. Fig. S2N). In dKO, Monocyte, Inflammatory, Restorative, MHC II but not Proliferating subsets were identified; interestingly, a unique subpopulation with highly induced CCL family genes (*Ccl7*, *Ccl4*, *Ccl2*, *Ccl8*, *Ccl6*, *Ccl24*, *Ccl3*) was detected and named CCL+ MPs (Fig. 2M and Suppl. Fig. S2O). When the pseudotime trajectory was analyzed, we found that in Ctrl (Fig. 2L) Monocytes plotted tightly together at the initial position of the trajectory line, followed by bifurcation into Restorative and Proliferating MPs. Inflammatory and MHC II MPs were distributed along the line between Monocyte and Restorative MPs. A distinct trajectory was plotted in dKO (Fig. 2M): starting from Monocyte, the line diverged into three branches, Restorative, MHC II, CCL+; Inflammatory mainly located along the two lines between Monocyte and Restorative/CCL+. The proportions of both Inflammatory and Restorative were increased in dKO vs. Ctrl (23.6% vs. 22.3%, 38.2% vs. 26.4%), but the MHC II MPs showed a decline (14.3% vs. 19.2%) (Fig. 2L-M); the dKO specific CCL+ cells displayed enhanced inflammatory features (enriched for *Rsad2*, *Ifit1* and *Tnf*) while the Ctrl specific Proliferating subset exhibited neutrophil-like scavenger characteristics (enriched for *S100a8*, *Camp* and *Ngp*). Altogether, the above results support that the inflammatory niche is skewed in dKO muscle. Consequently, by comparing the inflammatory score (defined by inflammatory gene set, Supp. Table S2), a significant elevation was detected (0.16 vs. –0.08) (Fig. 2N). Altogether, scRNA-seq results show that the intrinsic deletion of YY1 in MuSCs leads to remarkable niche remodeling primarily by altered FAP and MP compositions.

### Intrinsic deletion of YY1 from MuSCs enhances their crosstalk with MPs via CCL5/CCR5 axis

To delineate the molecular mechanism underlying the cellular composition changes in dKO muscles, bulk RNA-seq was performed on freshly isolated MuSCs from six pairs of Ctrl or dKO mice (Fig. 3A, Suppl. Table S3). A total of 1090 genes were up-regulated and 1527 down-regulated in dKO compared with Ctrl (Fig. 3B and Suppl. Table S3). Strikingly, Gene Ontology (GO) analysis revealed that the up-regulated genes were highly enriched for immune related terms such as “immune system process”, “innate immune response” etc. (Fig. 3C and Suppl. Table S3), suggesting an immune-like nature of dKO cells. Conversely, genes associated with “cell cycle” and “cell division” were predominantly down-regulated (Fig. 3D and Suppl. Table S3), aligning with the detected reduced proliferative capacity of dKO MuSCs (Suppl. Fig. S1D-E and Suppl. Table S3). A detailed examination of the up-regulated genes highlighted the induction of numerous pro-inflammatory mediators, particularly those facilitating MPs infiltration into injured tissue, such as a panel of chemoattractants from CCL family, *Ccl5*, *Ccl25*, *Ccl2*, *Ccl7*, *Ccl3* (Suppl. Fig. S3A-B and Suppl. Table S3) (*17, 32, 33*). Among these factors, *Ccl5* gene induction was robustly detected in dKO MuSCs (Fig. 3E, 2.58-fold) accompanied by the protein induction (Fig. 3F) and secretion (Fig. 3G); a gradually increased amount of CCL5 protein was also detected in dKO muscles after TMX injection (Fig. 3H). Furthermore, we found an escalated amount of *Ccl5* and *Ccr5* expression in isolated dKO MPs (Fig. 3I) and muscles (Fig. 3J). Co-localization of CCL5 and CCR5 proteins was also detected in dKO muscles (Fig. 3K). Altogether, these findings led us to hypothesize that CCL5 induction in dKO MuSCs enhances MP recruitment via a CCL5/CCR5 axis. To test this, transwell assay was performed by seeding MuSCs from Ctrl or dKO underneath the transwell insert and bone marrow derived macrophages (BMDMs) on the top (Fig. 3L). As expected, a much higher number of migrated BMDMs were detected in dKO vs. Ctrl (15.4 vs. 7.9) (Fig. 3L), indeed confirming the enhanced recruiting ability of dKO MuSCs. To validate the role of CCL5/CCR5 axis in mediating the recruitment, we found that BMDM attraction by dKO cells was significantly attenuated by neutralization of CCL5 (Fig. 3M) or CCR5 (Fig. 3N) with antibodies. Altogether, the above findings demonstrate that intrinsic loss of YY1 transforms MuSCs into immune-secretory and endows its heightened ability to crosstalk and recruit MPs to the niche microenvironment via the CCL5/CCR5 axis.

**Fig. 3.**
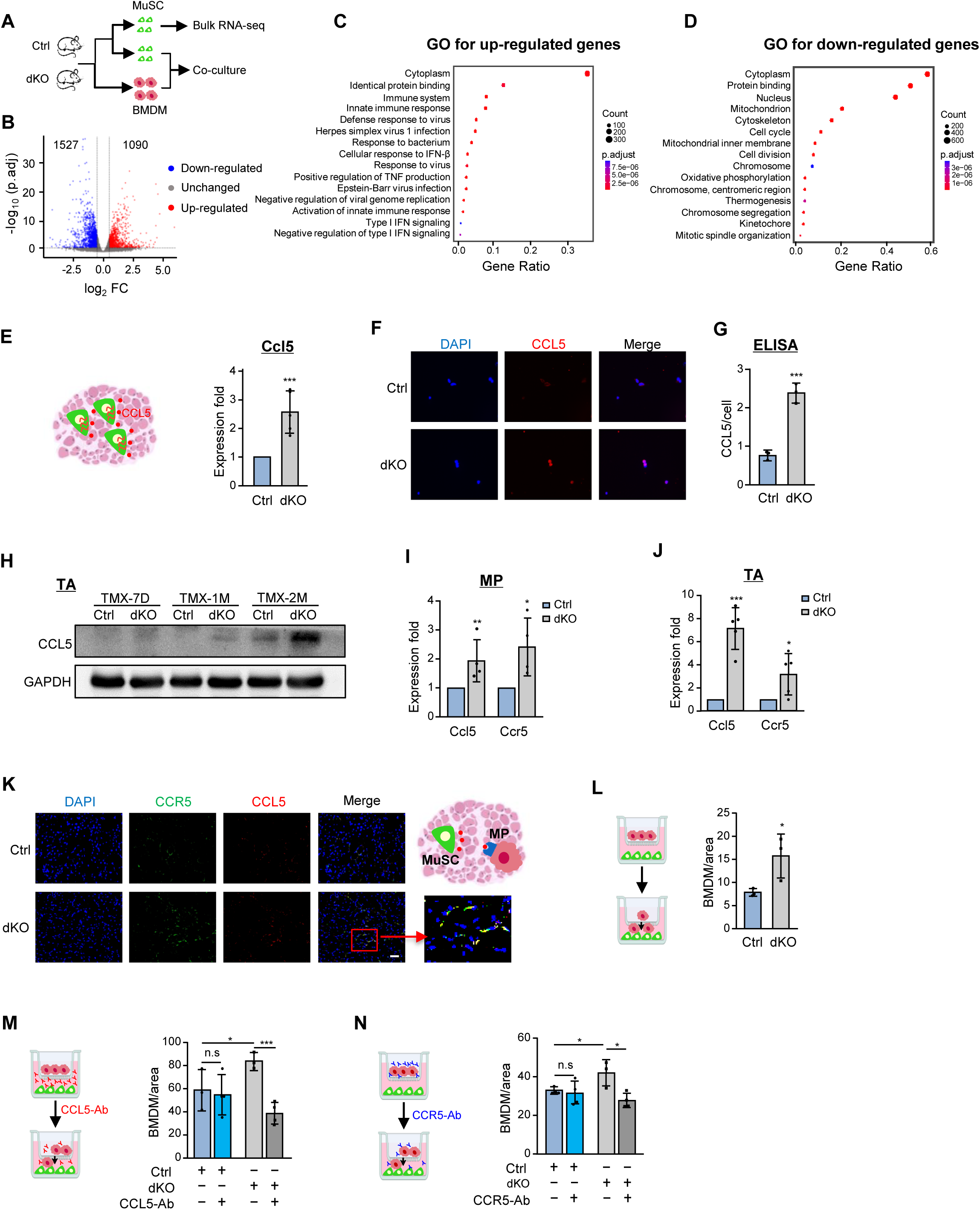
Intrinsic deletion of YY1 in MuSC induces enhanced crosstalk between MuSC and MP via CCL5/CCR5 axis. **A** Schematic of the experimental design for testing MuSC/MP interaction in Ctrl and dKO mice. **B** Differentiability expressed genes (DEGs) were identified from the RNA-seq profiling in Ctrl vs dKO MuSCs using Log_2_FC >0.5 as a cut-off. **C-D** GO analysis of the above identified 1090 up and 1527 down-regulated DEGs. **E** RT-qPCR detection of *Ccl5* mRNA in freshly isolated MuSCs form Ctrl and dKO. **F** IF staining of CCL5 protein in freshly isolated MuSCs from Ctrl and dKO. Scale bar: 25 μm, n=5. **G** ELISA detection of secreted CCL5 protein from Ctrl and dKO MuSCs, n=3. **H** Western blot detection of CCL5 protein in TA muscles of Ctrl and dKO at the designated times after TMX administration, n=3. **I** RT-qPCR detection of *Ccl5* and *Ccr5* mRNAs in freshly isolated MPs from Ctrl and dKO, n=4. **J** RT-qPCR detection of *Ccl5* and *Ccr5* mRNAs in Ctrl and dKO TA muscles. **K** IF staining of CCL5 and CCR5 proteins on TA muscles sections of Ctrl and dKO. Co-localization of CCL5 and CCR5 is shown in the red frame. Scale bar: 50 μm, n=5. **L** BMDMs were isolated from mdx mice and co-cultured with MuSCs from Ctrl and dKO in transwell. Quantification of migrated BMDMs is shown, n=3. **M-N** CCL5 or CCR5 antibody was added to the above transwell. Quantification of migrated BMDM is shown, n=3. All the bar graphs are presented as mean ±SD, Student’s t test (two-tailed unpaired) was used to calculate the statistical significance (E, G, I, J, L-N): *p < 0.05, **p < 0.01, ***p < 0.001, n.s. = no significance.

### TGFβ1 enrichment in the niche causes FAPaccumulation in dKO muscle by inhibiting apoptosis

To further elucidate the underlying cause for increased FAPs in dKO muscle, we examined if the dKO FAPs possess apoptosis resistance as suggested by Fig. 2K. PDGFRα+ FAPs were isolated from Ctrl and dKO mice and used for TUNEL staining (Fig. 4A). As expected, a significant reduction (32.5%) of TUNEL+ cells was observed in dKO vs. Ctrl (Fig. 4B), confirming the resistance to apoptosis of dKO FAPs. This was further supported by co-staining of TUNEL and PDGFRα on TA (Fig. 4C) or DP (Fig. 4D) muscles, showing a substantial decrease (9.3%, 26.3%) of apoptotic FAPs in dKO. Meanwhile, we also examined the proliferative ability of FAPs by EdU staining (Fig. 4A); no significant difference of the number of EdU+ cells was detected when the assay was conducted on cultured FAPs from Ctrl or dKO mice (Fig. 4E). However, when EdU incorporation was performed in vivo (Fig. 1A), a significant reduction (0.3% vs. 2.7%) of PDGFRα+ EdU+ FAPs was detected in dKO muscle (Fig. 4F), suggesting a potential decrease in proliferative capacity in the niche. Altogether, these results demonstrate that the accumulation of FAPs in dKO muscle probably arises from the mitigated apoptosis thus the enhanced survival.

**Fig. 4.**
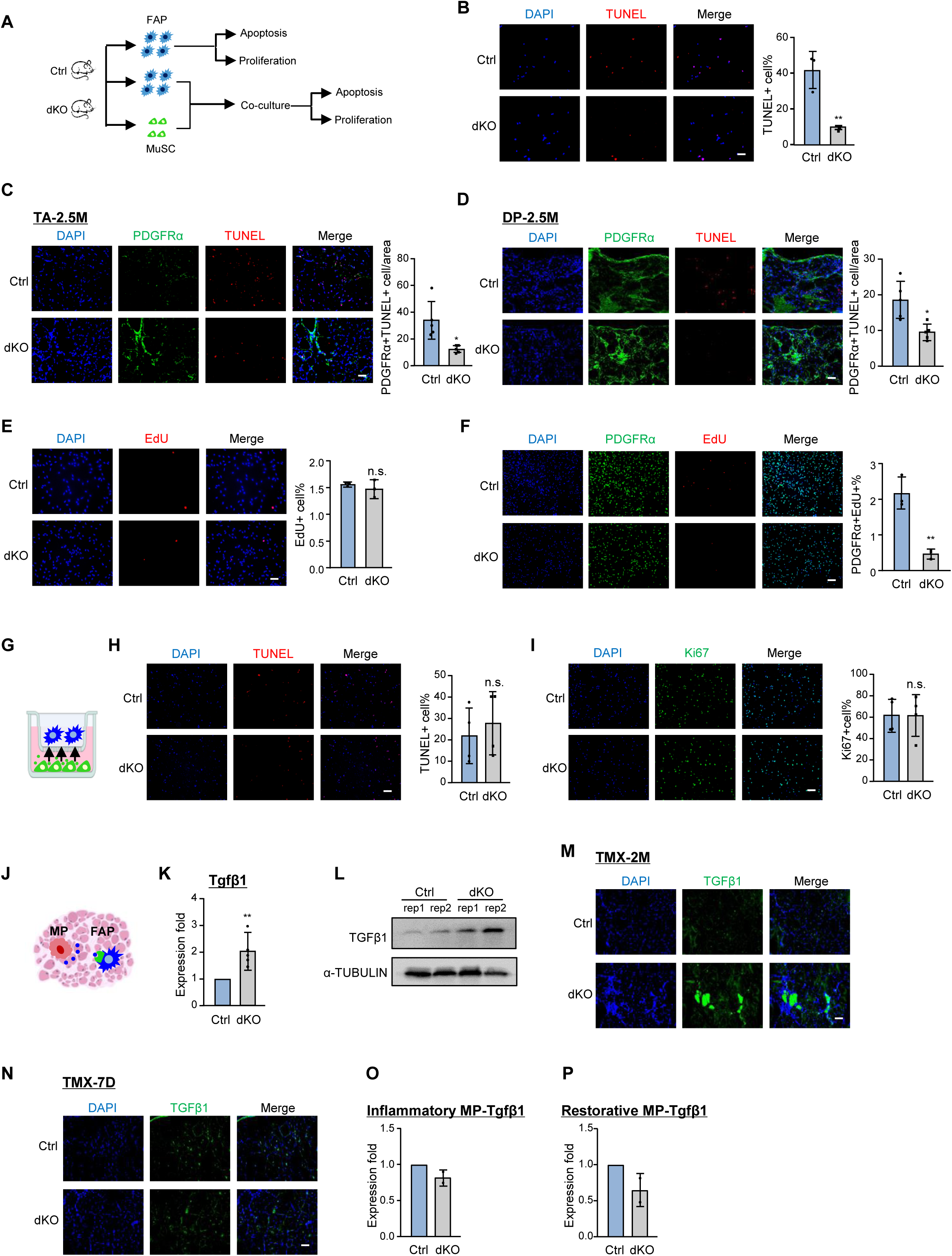
TGFβ1 enriched niche inhibits FAP apoptosis and causes FAP accumulation in dKO muscle. **A** Schematic of the experimental design for testing FAP/MuSC interaction in Ctrl and dKO mice. **B** TUNEL staining of FAPs isolated from Ctrl and dKO muscles. The percentage of TUNEL+ cells is shown. Scale bar: 50 μm, n=3. **C-D** TUNEL and PDGFRα staining of TA or DP muscles from Ctrl and dKO mice. The percentage of TUNEL+ PDGFRα+ cells is shown. Scale bar: 50 μm, n=5. **E** EdU staining of in vitro cultured (6 hr) FAPs from Ctrl and dKO. The percentage of EdU+ cells is shown. Scale bar: 50 μm, n=3. **F** EdU and PDGFRα staining of freshly isolated FAPs from EdU injected Ctrl and dKO mice (Fig. 1A). The percentage of EdU+ PDGFRα+ cells is shown. Scale bar: 100 μm, n=3. **G** Schematic of the MuSC and FAP co-culture experiment. **H** TUNEL staining of the above co-cultured FAPs. The percentage of TUNEL+ cells is shown Scale bar: 100 μm, n=4. **I** Ki67 staining of the above co-cultured FAPs. The percentage of Ki67+ cells is shown Scale bar: 100 μm, n=4. **J** Schematic of MP and FAP interaction. **K** RT-qPCR detection of TGFβ1 mRNA in TA muscles form Ctrl and dKO mice, n=5. **L** Western blot detection of TGFβ1 protein in TA muscles from Ctrl and dKO mice, n=2. **M-N** IF staining of TGFβ1 in TA muscles 2 months or 7 days after TMX administration. Scale bar: 50 μm, n=3. **O-P** RT-qPCR detection of TGFβ1 mRNA in inflammatory and restorative MPs macrophages isolated from Ctrl and dKO muscles, n=2. All the bar graphs are presented as mean ± SD, Student’s t test (two-tailed unpaired) was used to calculate the statistical significance (B-F, H-K, K): *p < 0.05, **p < 0.01, ***p < 0.001, n.s. = no significance.

To assess if direct crosstalk from MuSCs impacts FAP apoptosis and proliferation in dKO muscle niche (Fig. 4A), MuSCs from Ctrl or dKO mice were co-cultured with FAPs (isolated from Ctrl) for 24 hours (Fig. 4G) and there were no significant changes in apoptosis by TUNEL staining (Fig. 4H) or proliferation by Ki67 staining (Fig. 4I), suggesting that MuSCs may exert negligible direct impact on FAPs. We then sought to test the possibility that enhanced FAP survival in dKO muscle may be caused by increased TGFβ1 level since it was recently demonstrated that highly expressed TGFβ1 prevented FAP apoptosis and promoted their differentiation into matrix-producing cells, contributing to fibrosis in dystrophic muscle(*18*) (Fig. 4J). Indeed, much higher levels of *Tgfβ1*mRNA (Fig. 4K) and protein (Fig. 4L-M) were detected in dKO muscles at 2 months after YY1 deletion. Moreover, TGFβ1 signaling was barely detected 7 days after the YY1 deletion (TMX-7D) (Fig. 4N), indicating its gradual accumulation induced by the YY1 deletion in MuSCs. Since a quantity of studies suggested that MPs are the major contributor to the environmental TGFβ1 in dystrophic muscle niche(*18, 25, 34*), we further examined the TGFβ1 expression in MPs but found inflammatory (Fig. 4O) or restorative MPs (Fig. 4P) from dKO did not express increased levels, suggesting that the enriched TGFβ1 accumulation in dKO muscle niche may be a direct result of increased number of MPs. Altogether, these findings demonstrate that elevated TGFβ1 level inhibits FAP apoptosis and thus causes FAP accumulation to exacerbate fibrosis in dKO muscle.

### Inhibiting CCL5/CCR5 signaling axis with Maraviroc alleviates muscle dystrophy

To investigate if inhibiting above-defined CCL5/CCR5 signaling can mitigate dystrophy in dKO mice. To this end, we treated Ctrl and dKO mice with Maraviroc (MVC), a well-documented CCR5 antagonist(*35, 36*). Intraperitoneal administration of MVC in Ctrl and dKO mice was repeated bi-daily over a thirty-day period at 2 mg/kg after which the TA muscles were isolated for subsequent analysis (Fig. 5A). The treatment remarkably ameliorated the muscle phenotypes in dKO mice: the regeneration was evidently elevated (Fig. 5B), accompanied by significantly tampered amount of fibrosis (Fig. 5C-D) and inflammation (Fig. 5E). Interestingly, even in Ctrl mice, the treatment led to slightly attenuated pathological fibrosis and inflammation (Fig. 5C-E). As expected, a concomitant decline of the numbers of FAPs (Fig. 5F) and MPs (Fig. 5G) was observed in the dKO muscle niche, but the number of MuSCs did not show any obvious increase (Fig. 5H).

**Fig. 5.**
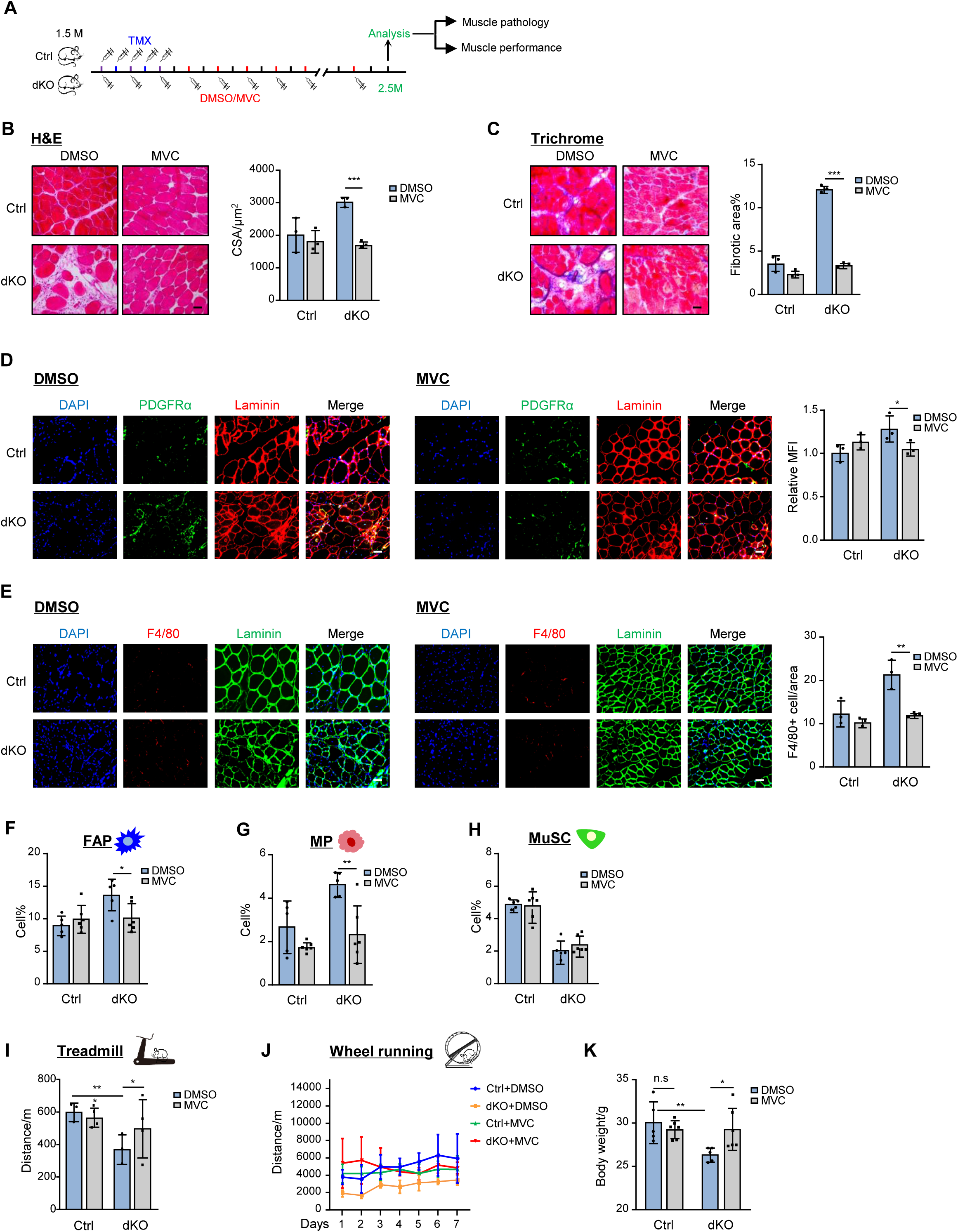
Targeting CCL5/CCR5 axis with MVC alleviates muscle dystrophy. **A** Schematic of the DMSO or MVC treatment and assessment in Ctrl and dKO mice. **B** Left: H&E staining of the TA and DP muscles collected after the treatment of Ctrl and dKO mice. Scale bar: 50 μm. Right: quantification of cross-sectional areas (CSAs) of fibers is shown, n≥3. **C** Left: Masson’s Trichrome staining of the above collected TA and DP muscles. Scale bar: 50 μm. Right: quantification of fibrotic areas is shown, n=3. **D** IF staining of DAPI (blue), PDGFRα (green) and Laminin (red) was performed on TA muscles of Ctrl and dKO mice after the treatment. The quantification of relative MFI of PDGFRα is shown. Scale bar: 50 μm, n=3. **E** IF staining of DAPI (blue), F4/80 (red) and Laminin (green) was performed on TA muscles of Ctrl and dKO mice after the treatment. The quantification of F4/80+ cell number per area is shown. Scale bar: 50 μm, n=3. **F-H** Flow cytometry detection of MP, FAP and MuSC population in Ctrl and dKO mice after the treatment, n=5. **I** Schematic of the muscle performance assessment by treadmill or voluntary wheel-running exercise. **J** The treated Ctrl and dKO mice were subjected to treadmill exercise and the running distance until exhaustion is shown, n=3. **K** The treated Ctrl and dKO mice were subjected to voluntary wheel-running exercise and the weekly running distance is shown, n=3. **L** Body weight of Ctrl and dKO mice after the treatment, n=5. All the bar graphs are presented as mean ±SD, Student’s t test (two-tailed unpaired) was used to calculate the statistical significance (C-I, K, M): *p < 0.05, **p < 0.01, ***p < 0.001, n.s. = no significance.

Consistently, the treated mice displayed significantly improved muscle function when subject to treadmill or voluntary wheel-running exercises. The endurance of dKO mice was notably recovered in the treadmill experiment (Fig. 5I). The treated dKO mice also demonstrated increased engagement in voluntary wheel-running, with a consistent increase in running distance over time (Fig. 5J). The overall morphology and health state of the dKO mice was improved by the treatment showing a significantly restored body weight (Fig. 5K). Altogether, these results further underpin the importance of CCL5/CCR5 signaling axis in muscle dystrophy and also highlight the possibility of targeting this axis to modulate muscle niche and decelerate disease progression.

### YY1 controls *Ccl5* expression in MuSC via regulating 3D E-P loop interaction

To answer how intrinsic YY1 deletion in MuSCs induces *Ccl5* expression, YY1 ChIP-seq was employed to map YY1 bound genes in freshly isolated MuSCs from mdx mice (Fig. 6A). A total of 4681 YY1 binding sites were identified with a canonical YY1 binding motif “AANATGGC” (Fig. 6B and Suppl. Table S4). Over half of the binding occurred in promoter regions (59%) while 29% and 12% in gene body and intergenic regions (Fig. 6C). GO analysis revealed YY1 bound genes were enriched for a wide range of terms such as “histone modification” and “mRNA processing” etc. (Fig. 6D). Further integration with the RNA-seq identified DEGs (Fig. 3B) uncovered a total of 181 and 226 YY1 bound genes were up– and down-regulated in dKO vs. Ctrl MuSCs (Fig. 6E and Suppl. Table S4). GO analysis revealed that the up-regulated targets were highly enriched for “Nucleus”, “Nucleoplasm” and “DNA binding” etc. while the down-regulated targets were related to “Nucleus”, “Nucleoplasm” and “Cell cycle” etc. (Suppl. Fig. S4A-B and Suppl. Table S4). Next, we took a close examination of the *Ccl5* locus and found no YY1 binding at its promoter region, instead, a binding site was identified at a distal site which was a potential enhancer by analyzing our previously published H3K27ac ChIP-seq data(*37*) (Fig. 6F). Considering YY1 is emerging as a 3D chromatin organizer that can facilitate enhancer-promoter (E-P) looping(*38–40*), we tested if YY1 modulates *Ccl5* transcription via orchestrating E-P looping in MuSCs. To this end, we performed in situ Hi-C(*41*) to interrogate the 3D genome organization in Ctrl and dKO MuSCs. As a result, our result revealed a mild global reorganization in chromatin interactions in dKO vs. Ctrl cells. We found that YY1 knock-out in MuSCs had limited effect on global 3D genome organization as no difference in the contact frequency was detected between Ctrl and dKO at both short-and long-distance levels (Suppl. Fig. S4C). The YY1 knock-out did not alter the overall genome organization at compartment level either (Suppl. Fig. S4D), indicated by infrequent locus switching between A and B compartments, with only 2.8% B to A and 2.2% A to B switching (Fig. 6G). At the TAD level, however, we noticed an evident TAD boundary remodeling: 9393 new boundaries formed while 4033 out of 13416 remained unchanged (Fig. 6H); and a significant decline in the TAD size was detected in dKO vs. Ctrl (Fig. 6I). Additionally, a slight increase in the number of TADs was observed (9557 in dKO vs. 8965 in Ctrl) (Fig. 6J). Furthermore, at the looping level, we found the loop size remained largely unaltered (Fig. 6K) while a declined number of looping events occurred in dKO vs. Ctrl (1056 vs. 1753) (Fig. 6L). These results suggest that YY1 may play a role in regulating the 3D genome in MuSCs via looping disruption.

**Fig. 6.**
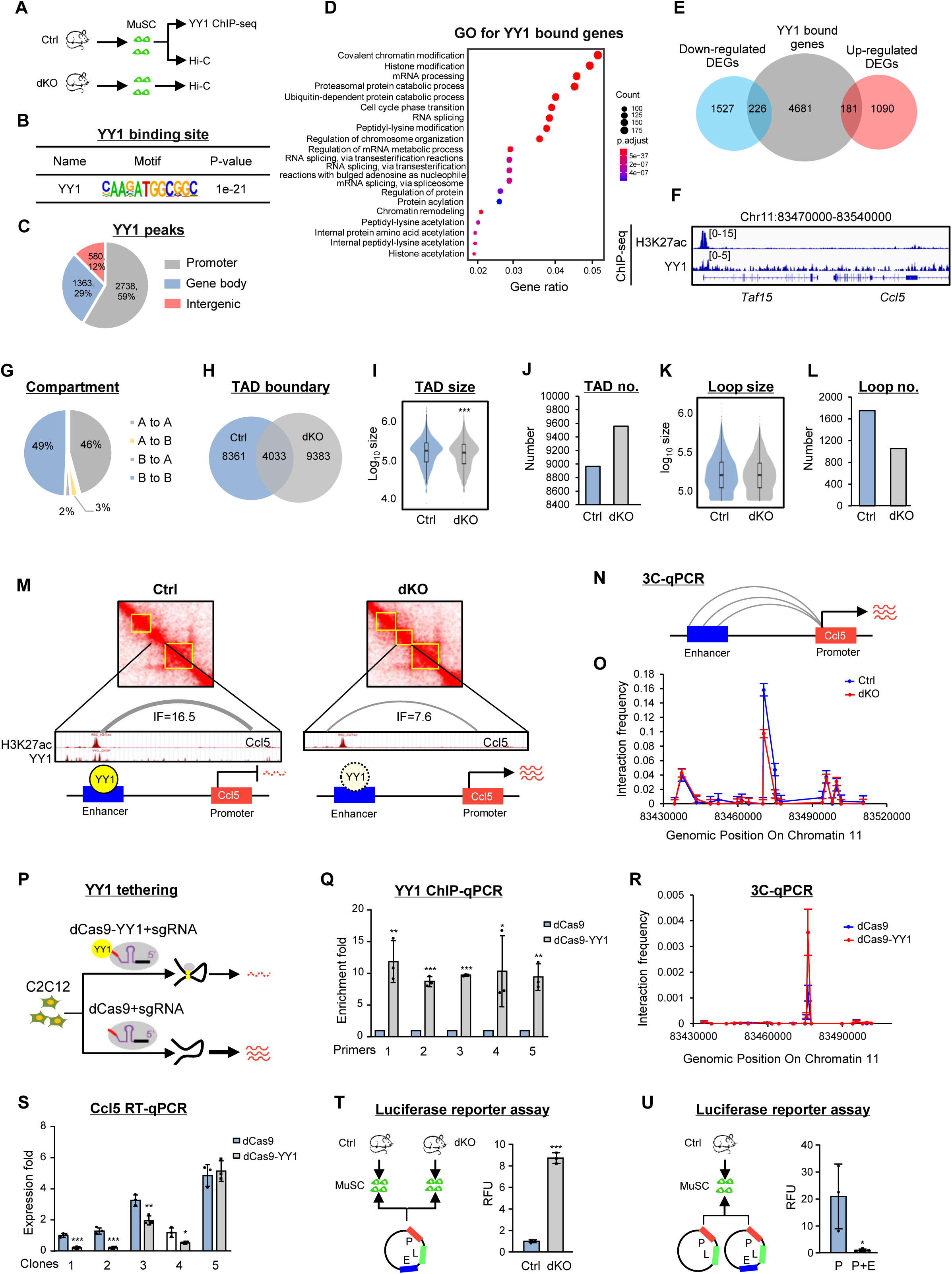
YY1 controls *Ccl5* expression in MuSC via regulating 3D looping interaction. **A** Schematic of the experimental design for dissecting YY1 regulation of *Ccl5* expression. **B** Enrichment of YY1 motif in the identified YY1 ChIP-seq binding regions. **C** Genomic distribution of the identified 4681 YY1 binding peaks. **D** GO analysis of the identified YY1 ChIP-seq targets. **E** Venn diagram showing the overlapping between the above identified YY1 ChIP-Seq targets and the down– or up-regulated genes. **F** Genomic snapshots on *Ccl5* locus showing co-binding of H3K27ac and YY1. **G** Pie chart showing HiC identified A/B compartment switch between Ctrl and dKO. **H** Venn diagram showing the changed TAD boundaries between Ctrl and dKO. **I-J** Comparison of TAD number and size between Ctrl and dKO. **K-L** Comparison of loop size and number comparison between Ctrl and dKO. **M** Heatmap showing the Hi-C interactions encompassing *Ccl5* locus (yellow box indicates TAD). Enhanced E-P interaction (grey curve) is shown in dKO vs Ctrl. **N** Schematic of 3C-qPCR experiment design. **O** 3C-qPCR detection of *Ccl5* enhancer proximal region on chromatin 11 showing decreased E-P interaction in dKO vs. Ctrl MuSC. **P** Schematic of YY1 tethering experiment design in C2C12 cells. **Q** ChIP-qPCR detection of YY1 enrichment on the tethered site in dCas9-YY1 vs. dCas9 cells, primers 1-5 were designed to target the proximal region of the tethered site, n=3. **R** 3C-qPCR detection of E-P interaction on *Ccl5* locus in dCas9-YY1 vs. dCas9 cells. **S** RT-qPCR detection of *Ccl5* expression in dCas9-YY1 vs. dCas9 cells, n=3. **T** Left: Luciferase reporter (L) with promoter (P) and enhancer (E) sequences was transfected into MuSCs from Ctrl and dKO. Right: Relative fluorescence unit (RFU) of reporter activity is shown, n=3. **U** Left: Luciferase reporters with *Ccl5* P or P+E sequences were transfected into Ctrl MuSC. Right: RFU of reporter activity is shown, n=3. All the bar graphs are presented as mean ±SD, Student’s t test (two-tailed unpaired) was used to calculate the statistical significance (I, K, Q, S-U): *p < 0.05, **p < 0.01, ***p < 0.001, n.s. = no significance.

Next, we took a close examination of the *Ccl5* locus, attempting to elucidate the cause behing high *Ccl5* in dKO MuSCs. Interestingly, two evident TADs were observed encompassing the *Ccl5* locus in Ctrl; and a newly gained third one was found in dKO and the above-identified YY1 bound enhancer was located within this gained TAD (Fig. 6M). Further analysis of E-P looping defined one strong looping event occurring between this enhancer and the *Ccl5* promoter; moreover, the interacting strength measured by interaction frequency was significantly lower in dKO vs. Ctrl (7.6 vs. 16.5) (Fig. 6M), suggesting an interesting scenario where YY1 mediated E-P looping functions to suppress *Ccl5* induction in Ctrl and the attenuation of the looping relieves the suppression in dKO. To test this notion, the above-identified E-P interaction was validated by Chromosome Conformation Capture (3C) assay in freshly isolated MuSCs from Ctrl and dKO using the *Ccl5* promoter as an anchor (Fig. 6N). Expectedly, a prominent interaction peak was detected at the enhancer region in both Ctrl and dKO groups and the quantified interaction frequency by 3C-qPCR weakened as the genomic distance from the enhancer site increased. In addition, the interaction frequency at the enhancer site was reduced in dKO vs. Ctrl (0.16 vs. 0.10) (Fig. 6O). These results indicating that the YY1 deletion in MuSCs indeed attenuates the E-P interaction at the *Ccl5* locus.

To further substantiate the direct function of YY1 in facilitating the E-P interaction, artificial tethering of YY1 protein to the *Ccl5* enhancer region (Fig. 6P) was performed in C2C12 myoblast cell line where no direct binding of YY1 on the *Ccl5* enhancer locus was detected by analyzing the existing H3K27ac and YY1 ChIP-seq data from C2C12 myoblast(*8, 42*) (Suppl. Fig. S4E). A dCas9-YY1 plasmid and five sgRNAs targeting the enhancer region were harnessed to generate five stable clones expressing increased level of dCas9-YY1; the transfection efficiency was confirmed in both mRNA (Suppl. Fig. S4F) and protein levels, with no significant influence on endogenous YY1 expression (Suppl. Fig. S4G). As a result, YY1 enrichment at the *Ccl5* enhancer region was markedly strengthened after tethering (Fig. 6Q) (detected by YY1 ChIP-qPCR using five primers targeting the enhancer region) and an increased contact frequency (0.0036 vs. 0.0012) was detected between the promoter and the enhancer by 3C-qPCR (Fig. 6R). Accordingly, the expression level of *Ccl5* was significantly reduced in four of the five stable clones (Fig. 6S). Altogether, the above results demonstrate an active role of YY1 in facilitating E-P interaction to curb the activation of *Ccl5* locus in Ctrl MuSCs. This conclusion was further corroborated by transfecting a luciferase reporter containing the promoter and enhancer regions of *Ccl5* in Ctrl and dKO MuSCs (Fig. 6T). A significantly higher level of luciferase activity was detected in dKO vs. Ctrl (1.00 vs. 8.73) (Fig. 6T). Moreover, a reporter devoid of the enhancer sequence displayed elevated reporter activity (20.86 vs 1.00) in Ctrl MuSCs (Fig. 6U). altogether, these results have confirmed the direct role of YY1 in repressing the *Ccl5* transcription by enhancing the E-P interaction.

## Discussion

In this study, we leverage the YY1/mdx dKO mouse strain to elucidate how MuSCs impact their niche microenvironment in dystrophic muscle. We demonstrate the MuSCs are capable of modulating their niche via cellular interactions with MPs and FAPs. Intrinsic deletion of YY1 in MuSCs leads to unexpected aggravation of inflammation and fibrosis in dystrophic mdx mice which is attributed to altered niche microenvironment characterized by skewed dynamics and heterogeneity of MPs and FAPs. Further investigation demonstrates that conveying the dKO’s intrinsic change to niche alterations stems from the induction of many immune related genes caused thus the conversion of MuSCs into competent secretory cells. In particular, we identify a CCL5/CCR5 signaling axis that mediates the MuSC-MP cellular interaction which enhances MP recruitment and exacerbates inflammation. As a result, elevated TGFβ leads to FAP accumulation and aggravated fibrosis. Treatment with the CCR5 antagonist MVC reduces dystrophic disease manifestations and improves muscle function. Furthermore, we also illuminate the intrinsic mechanism of how YY1 loss causes *Ccl5* induction in MuSCs via YY1’s ability to modulate enhancer-promoter looping interaction at *Ccl5* locus.

Our study provides compelling evidence to demonstrate the previously unappreciated active role of MuSCs in modulating niche microenvironment. As the small population of residents in skeletal muscle, MuSCs are often portrayed as the passive recipient of niche regulation. Mounting efforts are focused on dissecting how MuSC behavior is tightly regulated by spatiotemporal signaling from the niche and how niche-derived growth factors and signaling molecules, metabolic cues, the extracellular matrix and biomechanical cues, and immune signals exert their effects on MuSCs(*43*). However, sporadic evidence hinted the possible reciprocal effect that MuSCs exert on MPs. For example, a study from 2013 demonstrated that human myoblasts can indeed secrete an array of chemotactic factors to initiate monocyte chemotaxis in culture(*17*). A recent study from Oprescu et. al.(*28*) profiled transcriptomic dynamics at various stages of acute damage-induced muscle regeneration by scRNA-seq; a myoblast subpopulation enriched for immune genes were identified to be capable of active communication with immune cell populations. More recently, a study from Nakka K. et. al. demonstrates MuSCs can initiate the production of hyaluronic acid to modulate the ECM in the niche which in turn directs MuSCs to exist quiescence state for repair of injured muscle(*44*). Moreover, emerging reports reveal the presence of senescent MuSCs in regenerating and aging muscles; these senescent MuSCs are characterized by SASPs (senescence-associated secretory phenotypes) and could have paramount roles in remodeling niche microenvironment. Therefore, following our study, much effort will be needed to elucidate the functional interactions between MuSCs and the niche. Moreover, We also posit that similar niche-regulatory functions might be a feature of other adult stem cells; the tremendous advances in single cell profiling and spatial transcriptomic technologies are poised to revolutionize our understanding of stem cell niche dynamics in unparalleled detail.

Our findings uncovered a novel MuSC/MP interaction that offers insights into the pathogenic recruitment of MPs in dystrophic muscle. In acutely injured skeletal muscle, infiltrating MPs are mainly Ly6C^hi^ CCR2^+^CX3CR1^lo^ monocytes that are recruited through CCR2 chemotaxis signaling by myoblasts, endothelial cells and resident macrophages(*21*). These monocytes then differentiate into Ly6C^hi^ inflammatory MPs. After phagocytosing necrotic muscle debris, intramuscular Ly6C^hi^ MPs can switch into Ly6C^lo^ MPs. It has been shown that recruitment of Ly6^hi^ monocytes in mdx muscles is also dependent on CCR2. Nevertheless, the induction of other CC class chemokine ligands including CCL5, CCL6, CCL7, CCL8, and CCL9 and receptors CCR1, and CCR5, has also been observed in mdx muscle and suspected to have a role in MP recruitment(*45*). Here we for the first time demonstrate that MuSCs can also contribute to MP recruitment via the CCL5/CCR5 axis Nevertheless, MuSCs may not be the only source of CCL5. It is a known product of MPs, endothelial cells, eosinophils and also muscle fibers(*45*). Additionally, it will be interesting to further explore whether the MuSC-MP crosstalk has impacts on the heterogeneous presence of MPs in dystrophic muscles. It is also necessary to point out that mdx mice in fact display a milder phenotype compared to DMD patients. Muscle inflammation starts about 3 weeks of age, persist into 2-3 months of age, and then gradually subsides in limb muscles but not diaphragm. Our YY1/mdx dKO mice however display more severe dystrophic pathologies mimicking human DMD. Indeed, we found *Ccl5* expression was readily detectable in human DMD muscle tissues and displayed an inverse correlation with *YY1* mRNA level. Therefore, CCL5/CCR5 axis constitutes a potential target for DMD treatment. Although gene and molecular therapies targeting the primary defect of Dystrophin gene remain the most promising approach for DMD treatment, therapeutic strategies targeting the complex secondary mechanisms responsible for DMD pathogenesis are also being developed in parallel(*13*). Drugs aiming at reducing inflammation and fibrosis are proven to be effective in mitigating the disease progression. In our study, MVC treatment leads to remarkable amelioration of the dystrophy pathologies; both inflammation and fibrosis were reduced while muscle performance was improved, which encourages us to perform trials in human DMD patients in the future. In addition, our findings also reinforce the previously known pro-fibrotic role of TGFβ in dystrophic muscle highlighting it as a potential therapeutic target. Indeed, targeting TGFβ signaling by intramuscular injection of an inhibitor leads to reduced FAP accumulation and fibrosis in dystrophic mice(*34*). Altogether, our findings demonstrate that pharmacological drugs acting on the amelioration of the niche microenvironment can be good candidates for DMD treatment.

Lastly, in search of reasons underlying *Ccl5* induction upon YY1 deletion, we demonstrate that YY1 acts on *Ccl5* enhancer to orchestrate E-P interactions thus reinforcing its role as a structural protein. Emerging evidence demonstrates YY1 contributes to E-P structural interactions in a manner analogous to DNA interactions mediated by CTCF(*39*). It is shown that YY1 binds to active enhancers and promoter-proximal elements and forms dimers that facilitate the interaction of these DNA elements. Deletion of YY1 binding sites or depletion of YY1 protein disrupts E-P looping and gene expression(*39*). In line with this, reduced E-P interaction frequency on *Ccl5* locus was observed in dKO MuSCs which was reversed by artificially tethering YY1 protein to the *Ccl5* enhancer region. Interestingly, our results suggest that the YY1 facilitated E-P interaction suppresses but not enhances *Ccl5* induction. Studies from others and ours in fact also demonstrate that binding of certain TFs renders an enhancer element to become suppressive in gene expression(*39, 46, 47*). Additionally, we have only focused our investigation on *Ccl5*, it will also be interesting to elucidate how YY1 loss induces the expression of other *Ccl* family genes, for example, we observed YY1 binding at the promoters of *Ccl7* and *Ccl25* genes (data not shown), indicating a rather different mode of action of YY1 on these genes.

## Methods

### Mice

YY1^f/f^ and C57BL/10 ScSn DMDmdx (mdx) mouse strains were purchased from The Jackson Laboratory (Bar Harbor, ME, USA). The YY1-inducible conditional KO (YY1^iKO^) strain (Ctrl: Pax7^CreERT2/R26YFP^; YY1^+/+^, YY1^iKO^: Pax7^CreERT2/R26YFP^; YY1^f/f^ mice) was generated by crossing Pax7^CreERT2/R26YFP^ mice with YY1^f/f^ mice. The YY1/mdx double KO (YY1^dKO^) strain (Ctrl: Pax7^CreERT2/R26YFP^; YY1^+/+^; mdx, YY1^dKO^: Pax7^CreERT2/R26YFP^; YY1^f/f^; mdx) was generated by crossing YY1^iKO^ with mdx mice. Primers used for genotyping are shown in Suppl. Table S1.

### Animal procedures

Inducible deletion of YY1 was administrated by intraperitoneal (IP) injection of tamoxifen (TMX) (Sigma-Aldrich, T5648) at 100 mg/kg (body weight). For EdU (Sigma-Aldrich, 900584-50MG) incorporation assay in vivo, 5 mg EdU (diluted in 100 μl PBS) injection via IP per day was performed for 3 consecutive days, followed by FACS isolation of MuSCs or FAPs 12 hours later. Maraviroc (Sigma-Aldrich, PZ0002-25MG) treatment was administrated by IP injection at 20 mg/kg every two days for 30 days. For voluntary wheel-running test, mice were housed individually for 7 days in polycarbonate cages with 12-cm-diameter wheels equipped with optical rotation sensors (Yuyan Instrument, ARW). For treadmill test, mice were adapted to a treadmill (Panlab, Harvard Apparatus, 76-0895) with a 5°incline at an initial speed of 10 cm/s, followed by a stepwise increase of 5 cm/s every two min until their exhaustion.

### Cell lines and cell culture

Mouse C2C12 myoblast cells (CRL-1772) and 293T cells (CRL-3216) were obtained from American Type Culture Collection (ATCC) and cultured in DMEM medium (Gibco, 12800-017) with 10% fetal bovine serum (Invitrogen, 16000044), 100 units/ml of penicillin and 100 μg of streptomycin (P/S, Gibco,15140-122) at 37 °C in 5% CO_2_. All cell lines were tested negative for mycoplasma contamination.

### Fluorescence-activated cell sorting and culturing

Muscle stem cells, fibro-adipogenic progenitors and macrophages were sorted based on established method(*37, 48–53*). Briefly, entire hindlimb muscles from mice were digested with collagenase II (LS004177, Worthington, 1000 units per 1 ml) for 90 min at 37 °C, the digested muscles were then washed in washing medium (Ham’s F-10 medium (N6635, Sigma) containing 10% horse serum, heat-inactivated (HIHS, 26050088, Gibco, 1% P/S) before SCs were liberated by treating with Collagenase II (100 units per 1 ml) and Dispase (17105-041, Gibco, 1.1 unit per 1 ml) for 30 min. The suspensions were passed through a 20G needle to release cells. Mononuclear cells were filtered with a 40 μm cell strainer and sorted by BD FACSAria IV with the selection of the GFP+ (MuSCs); FITC-(CD45-, CD31-, ITGA7-) APC+(SCA1+) (FAPs); FITC-(Cd45-) APC-(Ly6G-) eFluor450+(CD11b+) (MPs). Flowjo V10.8.1 was used for analysis of flow cytometry data. MuSCs were cultured in Ham’s F10 medium with 20% FBS, 5ng/ml β-FGF (PHG0026, Thermo Fisher Scientific) and 1% P/S, on coverslips and culture wells which were coated with poly-D-lysine solution (p0899, Sigma) at 37 °C overnight and then coated with extracellular matrix (ECM) (E-1270, Sigma) at 4 °C for at least 6 h. FAPs and MPs were cultured in DMEM medium with 10% FBS and 1% P/S.

### Isolation of bone marrow derived macrophages (BMDMs)

Isolating BMDM was performed according to literature(*25, 54*) with slight modification. Briefly, bone marrow from adult mdx mouse was obtained by flushing femur and tibiae with DMEM and cells were cultured in DMEM containing 20% FBS and 20 ng/mL M-CSF (Thermo, PMC2044) for 6-7 days. The culture DMEM medium was changed on the third day. After 7 days of culturing, cells were carefully washed by PBS twice and collected for experiments.

### Cell proliferation and apoptosis analyses

For EdU incorporation assay, freshly sorted MuSCs were seeded into prepared coverslips and EdU was added to the culture medium for 6 hours before harvest. EdU staining was performed following the instruction of Click-iT® Plus EdU Alexa Fluor® 594 Imaging Kit (C10639, Thermo Fisher Scientific). Growing cells on coverslips were incubated with 10 μM EdU for a designated time before the fixation with 4% PFA for 20 min. EdU-labeled cells were visualized using “click” chemistry with an Alexa Fluor® 594-conjugated azide and cell nuclei were stained by DAPI (Life Technologies, P36931). Fluorescent images were captured with a fluorescence microscope (Leica). TUNEL (Terminal deoxynucleotidyl transferase dUTP nick end labeling) assay was performed following our previous publication(*4, 55*) and the instruction of In Situ Cell Death Detection Kit TMR red (Roche, 12156792910). Cells were cultured on coverslips for 36 h, followed by washing twice with PBS and fixing with 4% PFA for 15 min. TUNEL staining was carried out by adding reaction mixture of label solution for 30 min at dark. The coverslips were mounted with DAPI solution to stain the cell nucleus. Fluorescent images were captured with a fluorescence microscope (Leica).

### Co-culture assay

BMDMs or FAPs were re-suspended at the appropriate concentration in DMEM culture medium and seeded in the 24-well inserts (Corning, 353097, 353095). The bottom chamber was coated and seeded with or without MuSCs in GM one day before the seeding of BMDMs or FAPs to avoid the MuSCs adhering to the upper inserts. After seeding and assembling, the co-cultured cells were incubated at 37℃5% CO_2_ for 12-24 hours and then inserts were harvested for experiments.

### Plasmids

For constructing the dCas9-YY1 plasmid, pAW91.dCas9 (Addgene, 104372) and pAW90.dCas9-YY1 (Addgene, 104373) plasmids were purchased from Addgene. Five sgRNAs were designed by CRISPOR(*56*) to target the *Ccl5* upstream enhancer region and cloned into a lentiguide-puro vector at the BsmbI site(lentiguide-puro-sgRNA1,2,3,4,5). For constructing the *Ccl5*-enhancer/promoter luciferase report plasmid, a 2456 bp DNA fragment of *Ccl5* enhancer region and 1500 bp *Ccl5* promoter were cloned into a pGL3-basic (purchased from Promega) vector between MluI and HindIII sites.

### Luciferase report assay

MuSCs were co-transfected with the *Ccl5*-enhancer/promoter luciferase reporter plasmid and internal control Renilla reporter plasmid. Cells were harvested 48 hours after transfection through adding 100 μl lysis buffer and gently shaking for 15 min at room temperature. Luciferase activity was measured by the Dual-Luciferase kit (Promega, E1910) according to our previous publication(*37, 55*). The luminescent signal was recorded by Elmer VICTOR ^TM^ X multilabel reader. The ratio of the reporter signal and the Renilla control signal was compared between different samples for further analysis.

### dCas9-YY1 tethering

For virus production, HEK293T cells were grown to 50%–75% confluency on a 15 cm dish and then transfected with 15 μg of pAW91.dCas9 or pAW90.dCas9-YY1, 11.25 μg psPAX2 (Addgene, 12260), and 3.75 μg pMD2.G (Addgene, 12259). Viral supernatant was cleared of cells by filtering with 0.2 μm filter membrane (Pall Corporation 4612). 5 mL of vital supernatant was mixed with 5mL DMEM medium and added to C2C12 cells in the presence of polybrene (Santa Cruz Biotechnology, sc-134220) at 8 μg/mL. After 24 hours, viral media was removed and fresh media containing blasticidin (Thermo R21001) at 10 μg/mL. Cells were selected until all cells on non-transduced plates died, to obtain the blasticidin+ dCas9 C2C12 and dCas9-YY1 C2C12 cell lines. The tethering guide RNAs were packaged by the virus as described above using the lentiguide-puro-sgRNA1,2,3,4,5 plasmids and were transduced into dCas9 C2C12 and dCas9-YY1 C2C12 cells respectively. After 24 hours, viral media was removed and fresh media containing puromycin (Thermo A1113802) at 2.5 mg/mL. Cells were selected until all cells on non-transduced plates died. Double-positive cells (blasticidin+ puromycin+) were identified and expanded.

### RNA extraction and real-time PCR

Total RNAs were extracted using TRIzol reagent (Invitrogen) following the manufacturer’s protocol. For quantitative RT-PCR, cDNAs were reverse transcribed using HiScript III First-Strand cDNA Synthesis Kit (Vazyme, R312-01). Real-time PCR reactions were performed on a LightCycler 480 Instrument II (Roche Life Science) using Luna Universal qPCR Master Mix (NEB, M3003L). Sequences of all primers used can be found in Suppl. Table S1.

### Immunoblotting, immunofluorescence, and immunohistochemistry

For Western blot assays, according to our prior publication(*57–59*), cultured cells were washed with ice-cold PBS and lysed in cell lysis buffer. Whole cell lysates were subjected to SDS–PAGE and protein expression was visualized using an enhanced chemiluminescence detection system (GE Healthcare, Little Chalfont, UK) as described before(*60*). The following dilutions were used for each antibody: YY1 (Abcam ab109237; 1:1000), CCL5 (Abcam ab189841; 1:500), CCR5 (Abcam ab65850; 1:500), TGF-β1 (Abcam ab92486; 1:1000), α-TUBULIN (Santa Cruz Biotechnology sc-23948; 1:5000), GAPDH (Sigma-Aldrich G9545-100UL; 1:5000), CAS9 (CST 14697T; 1:1000). For immunofluorescence staining, cultured cells were fixed in 4% PFA for 15 min and blocked with 3% BSA within 1 hour. Primary antibodies were applied to samples with indicated dilution below and the samples were kept at 4°C overnight. For immunofluorescence staining(*4, 53, 59*), cultured cells or myofibers were fixed in 4% PFA for 15 min and permeabilized with 0.5% NP-40 for 10 min. Then cells were blocked in 3% BSA for 1 h followed by incubating with primary antibodies overnight at 4°C and secondary antibodies for one hour at RT. Finally, the cells were mounted with DAPI to stain the cell nucleus and images were captured by a Leica fluorescence microscope. Primary antibodies and dilutions were used as following PAX7 (Developmental Studies Hybridoma Bank; 1:50), YY1 (Abcam ab109237, 1:200), CCL5 (Abcam ab189841; 1:200), F4/80 (Abcam ab6640, 1:200), PDGFRα (R&D BAF1062; 1:200), Ki67 (Santa Cruz Biotechnology, sc-23900; 1:200). For immunohistochemistry(*4, 50, 53*), in brief, slides were fixed with 4% PFA for 15 min at room temperature and permeabilized in ice-cold methanol for 6 min at −20°C.Heat-mediated antigen retrieval with a 0.01 M citric acid (pH 6.0) was performed for 5 min in a microwave. After 4% BSA (4% IgG-free BSA in PBS; Jackson, 001-000-162) blocking, the sections were further blocked with unconjugated AffiniPure Fab Fragment (1:100 in PBS; Jackson, 115-007-003) for 30 min. The biotin-conjugated anti-mouse IgG (1:500 in 4% BBBSA, Jackson, 115-065-205) and Cy3-Streptavidin (1:1250 in 4% BBBSA, Jackson, 016-160-084) were used as secondary antibodies. Primary antibodies and dilutions were used as follows: CCL5 (Abcam ab189841; 1:200), CCR5 (Abcam ab65850; 1:200), TGF-β1 (Abcam ab92486; 1:200), PDGFRα (R&D BAF1062; 1:200), Collagen Ia1(Novus NBP1-30054; 1:200), F4/80 (Abcam ab6640), CD68 (Biorad MCA1957GA; 1:200), CD206 (Abcam ab64693;1:200), Laminin (Sigma-Aldrich L9393-100UL, 1:800), Ki67 (Santa Cruz Biotechnology SC-23900; 1:200); PAX7 (Developmental Studies Hybridoma Bank; 1:50), eMyHC (Leica NCL-MHC-d; 1:200) for staining of muscle cryosections. Images were slightly modified with ImageJ in which background was reduced using background subtraction and brightness and contrast were adjusted. H&E (Hematoxylin and eosin),was performed as previously described(*4, 60, 61*). Masson’s trichrome staining was performed according to the manufacturer’s (ScyTek Laboratories, Logan, UT) instructions.

### Single-cell RNA-seq and data analysis

Single-cell RNA-seq was performed on 10x genomics platform. Briefly, whole muscle cells were isolated with an additional step of viability validation by Propidium Iodide (PI) staining. Red blood cells were eliminated by ACK buffers (150M NH_4_Cl, 100mM KHCO_3_, 10mM EDTA-2Na) before sorting. After sorting, live cells were washed with 0.04% BSA in PBS twice and resuspended in the BSA solution at an appropriate concentration (300-1200 cells/μl). Suspended cells were counted under a microscope and Typan blue was used to examine the cell viability. After the confirmation of cell number and viability, library construction was performed following the manufacturer’s instructions for generation of Gel Bead-In Emulsions (GEMs) using the 10x Chromium system. The single cell RNA library was sequenced by Illumina HiSeq X Ten instrument. CellRanger 7.0.0 and Seurat 4.3.0 were used to analyze the single-cell data. Ctrl and dKO groups were initially merged together and filtered with quality control parameters (cells with more than 10% expression on mitochondrial genes, or fewer than 500 total features expressed were filtered out; Suppl. Fig. S2D-H). The two groups were integrated using FindIntegrationAnchors and IntegrateData function wrapped in Seurat package to minimize batch effects(*19, 28, 31*). Dimensionality reduction was performed through Principal Component Analysis (PCA). UMAP embedding parameters were based on the top 40 PCs and embedded in 2-dimensions to visualize the data. Cells from Ctrl and dKO groups were relatively evenly distributed in all clusters, indicating no major batch effect after treatment (Suppl. Fig. S2I). To annotate different cell types, differentially expressed genes among cell clusters were identified using FindAllMarkers function. Genes were identified as significantly differentially expressed if FDR < 0.05 and expression in at least 20% of cells. To dissect the subtypes of FAPs and macrophages, we conducted second-round clustering (sub-clustering) within each cell type. Differentially expressed gene markers were also examined for sub-clusters for manual subtype annotation. Monocle 2.22.0 was used for pseudotime trajectory analysis(*62*). The raw counts data from FAP and macrophage populations were used for trajectory inference and the top sub-cluster DEGs were used as input gene lists for trajectory construction analysis. AddModuleScore function wrapped in Seurat was used to calculate the average expression levels of each gene set of interest (Supp. Table S2) at single-cell level, yielding module scores named as anti-apoptotic, apoptotic score and inflammatory score.

### Bulk RNA-seq and data analysis

For RNA-seq (polyA+mRNA)(*50, 51*), total RNAs were subjected to polyA selection (Ambion, 61006) followed by library preparation using NEBNext Ultra II RNA Library Preparation Kit (NEB, E7770S). Libraries were paired-end sequenced with read lengths of 150 bp on Illumina HiSeq X Ten or Nova-seq instruments. The raw reads of RNA-seq were processed following the procedures described in our previous publication(*37*). Briefly, the adapter and low-quality sequences were trimmed from 3’ to 5’ ends for each read, and the reads shorter than 36 bp were discarded. The clean reads were aligned to mouse (mm10) reference genome with STAR. Next, Cufflinks was used to quantify the gene expression. Genes with an expression level change greater than 1.5-fold and a p-value of <0.01 were identified as DEGs between two stages/conditions. GO enrichment analysis was performed using R package clusterProfiler.

### ChIP-seq and data analysis

YY1 ChIP was performed following our previously described protocol(*4*). 10 μg of antibodies against YY1 (Santa Cruz Biotechnology, sc-1703), or normal mouse IgG (Santa Cruz Biotechnology, sc-2025) were used for immunoprecipitation. Immunoprecipitated genomic DNA was resuspended in 20 μl of water. For ChIP-seq DNA library construction, a NEBNext® Ultra™ II DNA Library Prep Kit for Illumina® (NEB, E7645S) was used according to the manufacturer’s instructions. Bioanalyzer analysis and qPCR were used to measure the quality of DNA libraries including the DNA size and purity. Finally, DNA libraries were sequenced on the Illumina Genome Analyzer II platform. The raw data were first pre-processed by initial quality assessment, adapter trimming, and low-quality filtering and then mapped to the mouse reference genome (mm10) using bowtie2(*63*), and only the non-redundant reads were kept. The protein DNA-binding peaks (sites) were identified using MACS293 with an input (IgG) sample as the background. During the peak calling, candidate peaks were compared with the background, dynamic programming was used to determine λ of Poisson distribution, and the P-value cutoff was set to 0.0001 for YY1 ChIP-Seq experiment.

### Hi-C and data analysis

Hi-C was performed according to previously described protocols(*37*). Libraries were prepared by on-bead reactions using the NEB Next Ultra II DNA Library Preparation Kit (NEB, E7645S). The beads were separated on a magnetic stand, and the supernatant was discarded. After washes, the beads were resuspended in 20 μl of 10 mM tris buffer and boiled at 98°C for 10 min. The elute was amplified for 10 to 13 cycles of PCR with Phanta Master Mix (Vazyme, P511-01), and the PCR products were purified using VAHTS DNA Clean Beads (Vazyme, N411-01). The Hi-C libraries were paired-end sequenced with read lengths of 150 bp on an Illumina HiSeq X Ten instrument. Data were analyzed by Hi-C Pro, juicer box software and mapping to mouse genome mm10. Raw Hi-C data were processed as previously described(*49, 64*). Briefly, the in-situ Hi-C data was processed with a standard pipeline HiC-Pro (*65*). First, adaptor sequences and poor-quality reads were removed using Trimmomatic (ILLUMINACLIP: TruSeq3-PE-2.fa:2:30:10; SLIDINGWINDOW: 4:15; MINLEN:50). The filtered reads were then aligned to mouse reference genome (mm10) in two steps: 1) global alignment was first conducted for all pair-end reads, 2) the unaligned reads were split into prospective fragments using restriction enzyme recognition sequence (GATCGATC) and aligned again. All aligned reads were then merged and assigned to restriction fragments, while low quality (MAPQ<30) or multiple alignment reads were discarded. Invalid fragments including unpaired fragments (singleton), juxtaposed fragments (re-ligation pairs), un-ligated fragments (dangling end), self-circularized fragments (self-cycle), and PCR duplicates were removed from each biological replicate. The remaining validated pairs from all replicates of each stage were then merged, followed by read depth normalization using HOMER (http://homer.ucsd.edu/homer/ngs/) and matrix balancing using iterative correction and eigenvector decomposition (ICE) normalization to obtain comparable interaction matrix between different stages.

Following previous procedure(*66*), to separate the genome into A and B compartments, the ICE normalized intra-chromosomal interaction matrices at 100-kb resolution were transformed to observe/expect contact matrices, and the background (expected) contact matrices were generated to eliminate bias caused by distance-dependent decay of interaction frequency and read depth difference(*37, 49*). Pearson correlation was then applied to the transformed matrices and the first principal component (PC1) of these matrices was divided into two clusters. The annotation of genes and the expression profile were used to assign positive PC1 value to gene-rich component as compartment A and negative PC1 value to gene-poor component as compartment B.

Normalized contact matrix at 10 kb resolution of each time point was used for TAD identification using TopDom(*67*). In brief, for each 10-kb bin across the genome, a signal of the average interaction frequency of all pairs of genome regions within a distinct window centered on this bin was calculated, thus TAD boundary was identified with local minimal signal within certain window. The falsely detected TADs without local interaction aggregation were filtered out by statistical testing. Invariant TADs were defined using following criteria: 1) the distance of both TAD boundaries between two conditions is no more than 10 kb; 2) the overlapping between two TADs should be larger than 80%; stage-specific TADs were defined otherwise. Loops are identified by using HiCCUPS module of Arrowhead with default parameter at 10 kb resolutions.

### Quantitative analysis of chromosome conformation capture assays (3C-qPCR)

3C-qPCR was performed following published protocols(*68*). The chromatin was cut by HindIII restriction enzyme to obtain DNA fragments. The promoter region of *Ccl5* was set as the anchor to detect the designed E-P interactions. Primers were designed targeting the nearby site of *Ccl5* enhancer region. 18s rRNA was used as an internal control for quantification normalization. Sequences of the oligos used in the study were included in Suppl. Table S1.

### Statistics and reproducibility

Data represent the average of at least three independent experiments or mice ± s.d. unless indicated. The statistical significance of experimental data was calculated by the Student’s t-test (two-sided). *p<0.05, **p<0.01, ***p<0.001 and n.s.: no significance (p≥0.05). The statistical significance for the assays conducted with MuSCs from the same mouse with different treatments was calculated by the student’s t-test (paired). *p < 0.05, **p < 0.01, ***p < 0.001, n.s. = no significance (p≥0.05). Specifically, a single zero-truncated negative binomial distribution was fit to the input data and each region was assigned a P value based on the fitted distribution. Representative images of at least three independent experiments are shown in Fig. 1D-L, O, P; 3F, K; 4B-F, H, I, M, N; 5C-F; and Supplementary Fig. S1B, C, J, K; 5A, E; 6D, E; S2A-C; S3F.

## Data availability

In situ Hi-C, ChIP-seq, bulk RNA-seq, and scRNA-seq data reported in this paper are deposited in the Gene Expression Omnibus database under accession GSE250204. All other data supporting the findings of this study are available from the corresponding author on reasonable request.

## Author contributions

Yang Li designed and performed most of experiments; Fengyuan Chen and Yeelo Cheung performed and helped with animal experiments; Chuhan Li and Qiang Sun analyzed RNA-seq, ChIP-seq, Hi-C and scRNA-seq data; Yu Zhao supervised Hi-C assay; Bénédicte Chazaud provided constructive suggestions and supervised co-culture experiments; Ting Xie contributed to the revision of manuscript; Hao Sun supervised computational analyses; Yang Li and Huating Wang wrote the manuscript, with inputs from all authors.

## Acknowledgements

This work was supported by National Key R&D Program of China to H.W. (project code: 2022YFA0806003); General Research Fund (GRF) from Research Grants Council (RGC) of the HongKong (HK) Special Administrative Region, China to H.W. (project codes: 14106521, 14100620, 14105823 and 14115319 to H.W.; 14105123, 14103522, 14120420 and 14120619 to H.S.); Theme-based Research Scheme (TRS) from RGC (project code:T13-602/21-N); Collaborative Research Fund (CRF) from the Research Grants Council (RGC) of the Hong Kong Special Administrative Region, China (project code: C6018-19GF); Health and Medical Research Fund (HMRF) from Health Bureau of HK to H.W. (project codes: 10210906 and 08190626); the National Natural Science Foundation of China (NSFC) to H.W. (project codes: 82172436 and 31871304); the research funds from Health@InnoHK program launched by Innovation Technology Commission, the Government of HK to H.W.; CUHK Strategic Seed Funding for Collaborative Research Scheme (SSFCRS) to H.W.; Area of Excellence Scheme (AoE) from RGC (project code: AoE/M-402/20).

## Inventory of Supplemental Information

### 1. Supplemental Figures

Suppl. Fig. S1. Inducible deletion of YY1 in MuSCs aggravates muscle dystrophy in mdx mouse.

Suppl. Fig. S2. Intrinsic YY1 deletion in MuSCs alters cellular microenvironment in dystrophic muscle.

Suppl. Fig. S3. Intrinsic deletion of YY1 in MuSC induces enhanced crosstalk between MuSC and MP via CCL5/CCR5 axis.

Suppl. Fig. S4. YY1 controls *Ccl5* expression in MuSC via regulating 3D looping interaction.

### 2. Supplemental Tables

Suppl. Table S1. Sequences of oligonucleotides used in the study

Suppl. Table S2. Single-cell RNA-seq profiling in Ctrl and YY1 dKO MuSCs

Suppl. Table S3. Bulk RNA-seq in YY1 dKO MuSCs

Suppl. Table S4. YY1 ChIP-seq in mdx MuSCs

Suppl. Table S5. Hi-C analysis in Ctrl and YY1 dKO MuSCs

### 3. Supplementary figure legends

**Suppl. Fig. S1.**
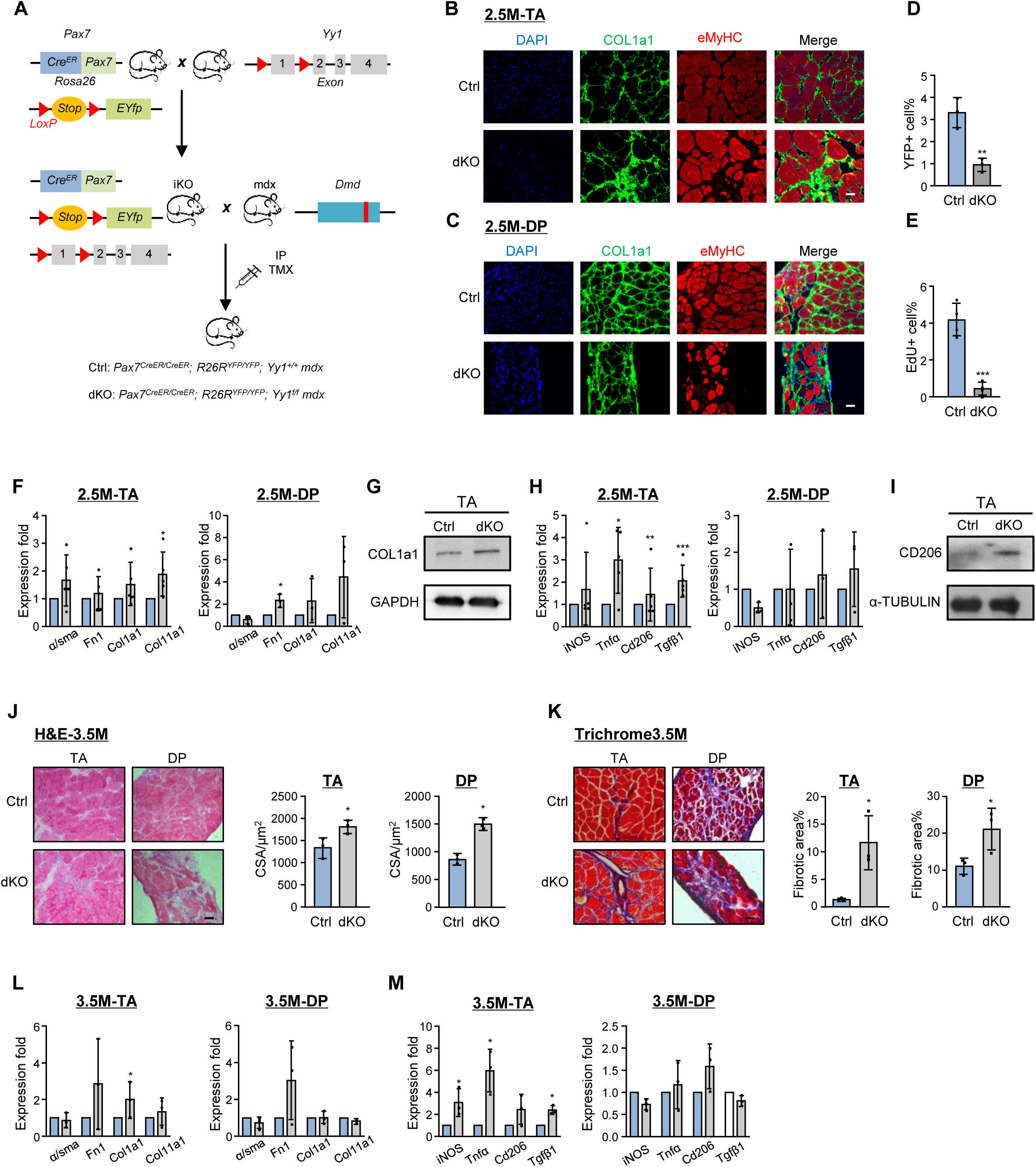
Inducible deletion of YY1 in MuSCs aggravates muscle dystrophy in mdx mouse. **A** Breeding scheme for generating inducible YY1 conditional knock out mdx (dKO: Pax7^CreERT2/R26YFP^; YY1^f/f^; mdx) and Control (Ctrl: Pax7^CreERT2/R26YFP^; YY1^+/+^; mdx) mice. **B-C** IF staining of DAPI (blue), COL1a1 (green) and eMyHC (red) was performed on the TA and DP muscles from 2.5M Ctrl and dKO mice. Scale bar: 50 μm. **D** MuSC pool was detected by flow cytometry (FC) with the YFP+ (Pax7+) signal, cells were freshly isolated from 2.5M Ctrl or dKO mice, n=3. **E** In vivo EdU incorporation was performed as Fig. 1A described, MuSCs were isolated for EdU+ cell quantification from 2M Ctrl or dKO mice, n=4. **F** RT-qPCR detection of fibrotic marker genes in TA and DP muscles from Ctrl and dKO mice, n=4. **G** Western blot detection of COL1a1 expression in TA muscles from 2.5 M Ctrl and dKO mice, n=3. **H** RT-qPCR detection of inflammatory marker genes in TA and DP muscles from 2.5M Ctrl and dKO mice. n=5. **I** Western blot detection of CD206 expression in TA muscles from 2.5M Ctrl and dKO mice, n=3. **J** Left: H&E staining of TA and DP muscles from 3.5 M Ctrl and dKO mice. Scale bar: 50 μm. Right: quantification of cross-sectional areas (CSAs) of the stained fibers, n=3. **K** Left: Masson’s Trichrome staining of the above muscles from. Scale bar: 50 μm. Right: quantification of the stained fibrotic areas, n=3. **L** RT-qPCR detection of fibrotic marker genes in TA and DP muscles from 3.5M Ctrl and dKO mice, n=3. **M** RT-qPCR detection of inflammatory marker genes in the above muscles, n=3. All the bar graphs are presented as mean ±SD, Student’s t test (two-tailed unpaired) was used to calculate the statistical significance (D-F, H, J-M): *p < 0.05, **p < 0.01, ***p < 0.001, n.s. = no significance.

**Suppl. Fig. S2.**
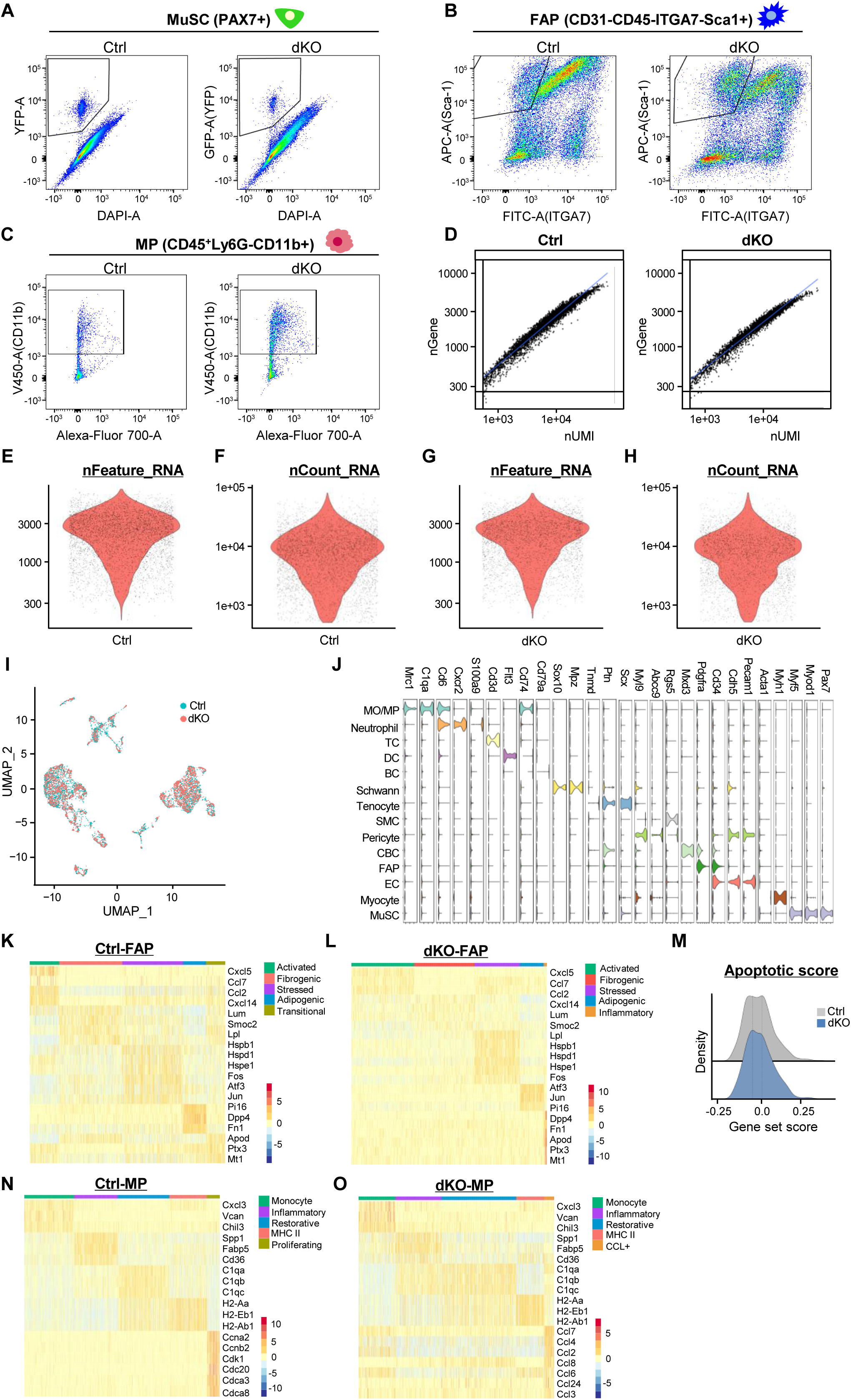
Intrinsic YY1 deletion in MuSCs alters cellular microenvironment in dystrophic muscle. **A-C** Representative FACS plots of isolating MuSCs (YFP+), FAPs (FITC-, APC+) and MPs (FITC-, APC-, eFluor450+). About 100,000 cells were sorted by FACS from Ctrl and dKO mice at 7, 21 and 60 days post-TMX injection. The percentage of cells is shown in Fig 2B-D. **D** Scatter plots showing the correlation between the number of genes detected (nGene, X-axis) and the number of RNA counts (nUMI, Y-axis). **E-H** Violin plots showing the distribution of the number of features or fraction counts of detected RNAs in Ctrl and dKO muscles. **I** Uniform manifold approximation projection (UMAP) embedding of single-cell data, colored by Ctrl or dKO. **J** Violin plots grouped by meta-clusters showing cell-type marker gene expression, which was used to classify clusters. **K-N** Heatmap of top 5 genes as determined by FindAllMarkers on the subclustered FAPs or MPs in Ctrl and dKO. **K** Ridge map showing the global distribution density of apoptotic score of FAPs.

**Suppl. Fig. S3.**
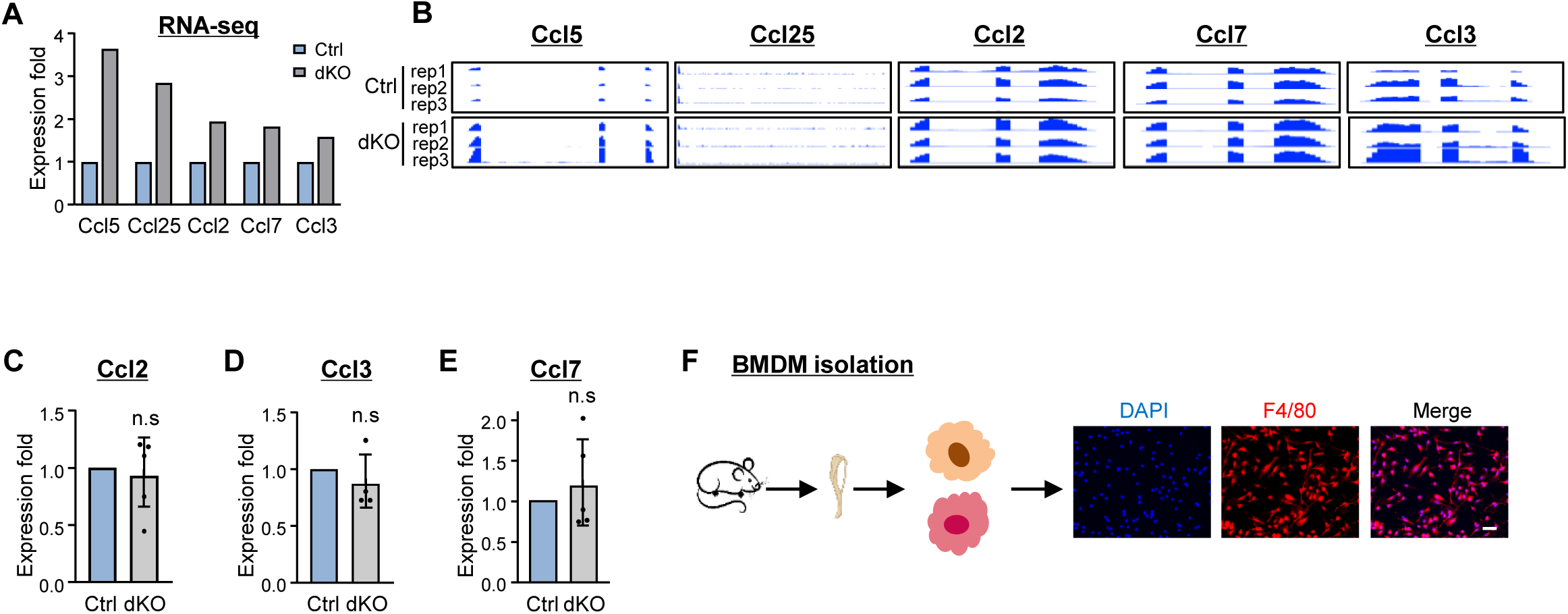
Intrinsic deletion of YY1 in MuSC induces enhanced crosstalk between MuSC and MP via CCL5/CCR5 axis. **A** Expression fold change of up-regulated *Ccl5*, *Ccl25*, *Ccl2*, *Ccl7*, *Ccl3* genes from RNA-seq data. **B** Genomic snapshots of the above *Ccl* genes. **C-E** RT–qPCR validation of the expression of *Ccl2*, *Ccl3*, *Ccl7* genes in Ctrl and dKO, n=5. **F** Schematic of Bone Marrow Derived Macrophage (BMDM) isolation from xxx and validation by F4/80 staining. Scale bar: 50 μm. All the bar graphs are presented as mean ±SD, Student’s t test (two-tailed unpaired) was used to calculate the statistical significance (C-E): *p < 0.05, **p < 0.01, ***p < 0.001, n.s. = no significance.

**Suppl. Fig. S4.**
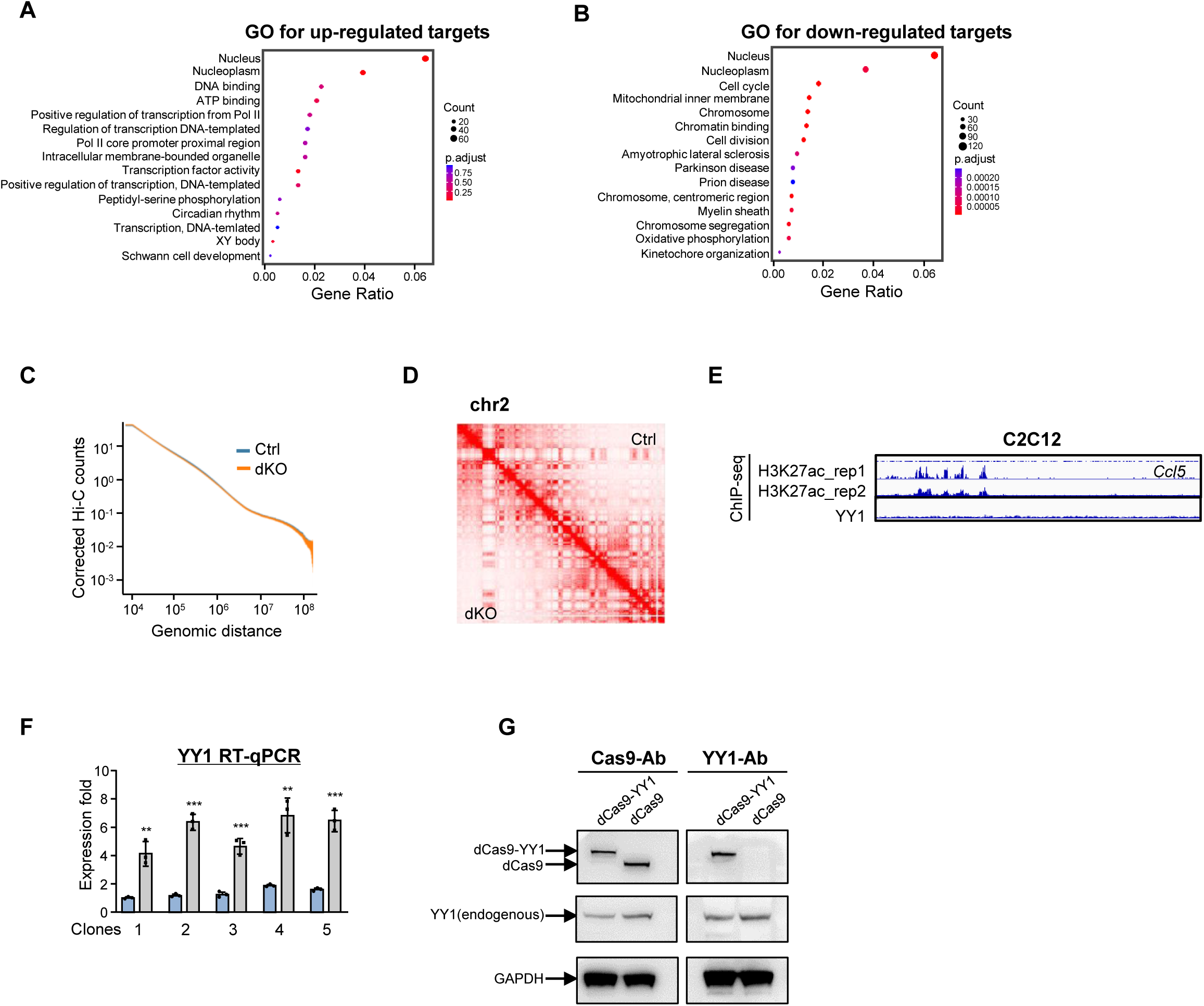
YY1 controls *Ccl5* expression in MuSC via regulating 3D looping interaction. **A-B** GO analysis of overlapped up– or down-regulated genes in Fig. 6E. **C** Contact frequency as a function of genomic distance along the whole genome in Ctrl and dKO MuSCs. **D** Representative heatmap showing the called compartment at chromosome 2 in Ctrl (upper triangle) and dKO (lower triangle) MuSCs. **E** Genomic snapshots of H3K27ac and YY1 ChIP-seq tracks in C2C12 showing no YY1 binding peak on *Ccl5* upstream enhancer sites. **F** RT-qPCR detection of overexpressed YY1 mRNAs in dCas9-YY1 vs. dCas9 transfected C2C12 cells, n=3. **G** Western blot validation of the expression of dCas9 or dCas9-YY1 in the above transfected C2C12 cells, the expression of endogenous YY1 protein is also shown. All the bar graphs are presented as mean ±SD, Student’s t test (two-tailed unpaired) was used to calculate the statistical significance (F): *p < 0.05, **p < 0.01, ***p < 0.001, n.s. = no significance.

## Notes

### Competing Interest Statement

The authors have declared no competing interest.

### Summary of Updates

update the mechanism figure

## References

1. A. Aziz, S. Sebastian, F. J. Dilworth, The origin and fate of muscle satellite cells. Stem Cell Rev 8, 609–622 (2012).

2. C. F. Bentzinger, Y. X. Wang, M. A. Rudnicki, Building muscle: molecular regulation of myogenesis. Cold Spring Harb Perspect Biol 4, (2012).

3. F. Relaix, P. S. Zammit, Satellite cells are essential for skeletal muscle regeneration: the cell on the edge returns centre stage. *Development (Cambridge*, England) 139, 2845–2856 (2012).

4. F. Chen et al., YY1 regulates skeletal muscle regeneration through controlling metabolic reprogramming of satellite cells. EMBO J 38, (2019).

5. L. Zhou et al., Linc-YY1 promotes myogenic differentiation and muscle regeneration through an interaction with the transcription factor YY1. Nat Commun 6, 10026 (2015).

6. K. Sun, L. Zhou, Y. Zhao, H. Wang, H. Sun, Genome-wide RNA-seq and ChIP-seq reveal Linc-YY1 function in regulating YY1/PRC2 activity during skeletal myogenesis. Genomics data 7, 247–249 %@ 2213-5960 (2016).

7. K. Sun, L. Lu, H. Wang, H. Sun, Genome-wide profiling of YY1 binding sites during skeletal myogenesis. Genomics Data 2, 89–91 (2014).

8. L. Lu et al., Genome-wide survey by ChIP-seq reveals YY1 regulation of lincRNAs in skeletal myogenesis. 32, 2575–2588 (2013).

9. K. Ji et al., Skeletal muscle increases FGF21 expression in mitochondrial disorders to compensate for energy metabolic insufficiency by activating the mTOR–YY1–PGC1α pathway. Free Radical Biology and Medicine 84, 161–170 (2015).

10. O. Mashinchian, A. Pisconti, E. Le Moal, C. F. Bentzinger, The Muscle Stem Cell Niche in Health and Disease. Curr Top Dev Biol 126, 23–65 (2018).

11. J. Farup, L. Madaro, P. L. Puri, U. R. Mikkelsen, Interactions between muscle stem cells, mesenchymal-derived cells and immune cells in muscle homeostasis, regeneration and disease. Cell Death Dis 6, e1830 (2015).

12. B. Biferali, D. Proietti, C. Mozzetta, L. Madaro, Fibro-Adipogenic Progenitors Cross-Talk in Skeletal Muscle: The Social Network. Front Physiol 10, 1074 (2019).

13. O. Cappellari, P. Mantuano, A. De Luca, “The Social Network” and Muscular Dystrophies: The Lesson Learnt about the Niche Environment as a Target for Therapeutic Strategies. Cells 9, (2020).

14. B. J. T. i. I. Chazaud, Inflammation and Skeletal Muscle Regeneration: Leave It to the Macrophages!, (2020).

15. M. Saclier, S. Cuvellier, M. Magnan, R. Mounier, B. Chazaud, Monocyte/macrophage interactions with myogenic precursor cells during skeletal muscle regeneration. FEBS J 280, 4118–4130 (2013).

16. J. Dort, P. Fabre, T. Molina, N. A. Dumont, Macrophages Are Key Regulators of Stem Cells during Skeletal Muscle Regeneration and Diseases. Stem Cells Int 2019, 4761427 (2019).

17. B. Chazaud et al., Satellite cells attract monocytes and use macrophages as a support to escape apoptosis and enhance muscle growth. J Cell Biol 163, 1133–1143 (2003).

18. D. R. Lemos et al., Nilotinib reduces muscle fibrosis in chronic muscle injury by promoting TNF-mediated apoptosis of fibro/adipogenic progenitors. Nat Med 21, 786–794 (2015).

19. K. K. Saleh et al., Single cell sequencing maps skeletal muscle cellular diversity as disease severity increases in dystrophic mouse models. Iscience 25, (2022).

20. D. D. Scripture-Adams et al., Single nuclei transcriptomics of muscle reveals intra-muscular cell dynamics linked to dystrophin loss and rescue. Communications biology 5, 989 (2022).

21. X. Wang, L. Zhou, The many roles of macrophages in skeletal muscle injury and repair. Frontiers in Cell and Developmental Biology 10, 952249 (2022).

22. P. Singh, B. Chazaud, Benefits and pathologies associated with the inflammatory response. Experimental Cell Research 409, 112905 (2021).

23. W. Chen, W. You, T. G. Valencak, T. Shan, Bidirectional roles of skeletal muscle fibro-adipogenic progenitors in homeostasis and disease. Ageing Research Reviews, 101682 (2022).

24. R. Farahzadi, B. Valipour, S. Montazersaheb, E. Fathi, Targeting the stem cell niche micro-environment as therapeutic strategies in aging. Frontiers in Cell and Developmental Biology 11, 1162136 (2023).

25. G. Juban et al., AMPK Activation Regulates LTBP4-Dependent TGF-beta1 Secretion by Pro-inflammatory Macrophages and Controls Fibrosis in Duchenne Muscular Dystrophy. Cell Rep 25, 2163–2176 e2166 (2018).

26. M. Low, C. Eisner, F. Rossi, Fibro/Adipogenic Progenitors (FAPs): Isolation by FACS and Culture. Methods Mol Biol 1556, 179–189 (2017).

27. L. Ertoz, M. Steinbach, V. Kumar, in Workshop on clustering high dimensional data and its applications at 2nd SIAM international conference on data mining. (2002), vol. 8.

28. S. N. Oprescu, F. Yue, J. Qiu, L. F. Brito, S. Kuang, Temporal Dynamics and Heterogeneity of Cell Populations during Skeletal Muscle Regeneration. iScience 23, 100993 (2020).

29. D. Kennedy, R. Jäger, D. D. Mosser, A. Samali, Regulation of apoptosis by heat shock proteins. IUBMB life 66, 327–338 (2014).

30. D. Lanneau et al., Heat shock proteins: essential proteins for apoptosis regulation. Journal of cellular and molecular medicine 12, 743–761 (2008).

31. A. J. De Micheli et al., Single-Cell Analysis of the Muscle Stem Cell Hierarchy Identifies Heterotypic Communication Signals Involved in Skeletal Muscle Regeneration. Cell Rep 30, 3583–3595 e3585 (2020).

32. L. Yahiaoui, D. Gvozdic, G. Danialou, M. Mack, B. J. Petrof, CC family chemokines directly regulate myoblast responses to skeletal muscle injury. J Physiol 586, 3991–4004 (2008).

33. N. Unver, Macrophage chemoattractants secreted by cancer cells: Sculptors of the tumor microenvironment and another crucial piece of the cancer secretome as a therapeutic target. Cytokine Growth Factor Rev 50, 13–18 (2019).

34. D. A. Mazala, et al., TGF-beta-driven muscle degeneration and failed regeneration underlie disease onset in a DMD mouse model. JCI Insight 5, (2020).

35. A. M. Passman et al., Maraviroc prevents hcc development by suppressing macrophages and the liver progenitor cell response in a murine chronic liver disease model. 13, 4935 (2021).

36. Z. Zeng, T. Lan, Y. Wei, X. J. G. Wei, diseases, CCL5/CCR5 axis in human diseases and related treatments. (2021).

37. Y. Zhao et al., Multiscale 3D genome reorganization during skeletal muscle stem cell lineage progression and aging. Science Advances 9, eabo1360 (2023).

38. L. Li et al., YY1 interacts with guanine quadruplexes to regulate DNA looping and gene expression. Nat Chem Biol 17, 161–168 (2021).

39. A. S. Weintraub et al., YY1 Is a Structural Regulator of Enhancer-Promoter Loops. Cell 171, 1573–1588 e1528 (2017).

40. J. A. Beagan et al., YY1 and CTCF orchestrate a 3D chromatin looping switch during early neural lineage commitment. Genome Res 27, 1139–1152 (2017).

41. S. S. Rao et al., A 3D map of the human genome at kilobase resolution reveals principles of chromatin looping. Cell 159, 1665–1680 (2014).

42. P. Asp et al., Genome-wide remodeling of the epigenetic landscape during myogenic differentiation. Proceedings of the National Academy of Sciences 108, E149–E158 (2011).

43. M. R. Hicks, A. D. Pyle, The emergence of the stem cell niche. Trends in cell biology, (2023).

44. K. Nakka et al., JMJD3 activated hyaluronan synthesis drives muscle regeneration in an inflammatory environment. Science 377, 666–669 (2022).

45. J. D. Porter et al., Persistent over-expression of specific CC class chemokines correlates with macrophage and T-cell recruitment in mdx skeletal muscle. Neuromuscular Disorders 13, 223–235 (2003).

46. X. L. Peng et al., MyoD-and FoxO3-mediated hotspot interaction orchestrates super-enhancer activity during myogenic differentiation. Nucleic acids research 45, 8785–8805 (2017).

47. P. Kolovos, T. A. Knoch, F. G. Grosveld, P. R. Cook, A. Papantonis, Enhancers and silencers: an integrated and simple model for their function. Epigenetics & chromatin 5, 1–8 (2012).

48. L. He, Z. He, Y. Li, H. Sun, H. Wang, in *Skeletal Muscle Stem Cells: Methods and Protocols*. (Springer, 2023), pp. 287–311.

49. L. He et al., CRISPR/Cas9/AAV9-mediated in vivo editing identifies MYC regulation of 3D genome in skeletal muscle stem cell. Stem Cell Reports 16, 2442–2458 (2021).

50. S. Zhang et al., ATF3 induction prevents precocious activation of skeletal muscle stem cell by regulating H2B expression. Nature Communications 14, 4978 (2023).

51. Z. He et al., Sugt1 loss in skeletal muscle stem cells impairs muscle regeneration and causes premature muscle aging. Life Medicine, lnad039 (2023).

52. Y. Li et al., Long noncoding RNA SAM promotes myoblast proliferation through stabilizing Sugt1 and facilitating kinetochore assembly. 11, 1–16 (2020).

53. X. Chen et al., Translational control by DHX36 binding to 5′ UTR G-quadruplex is essential for muscle stem-cell regenerative functions. Nature Communications 12, 5043 (2021).

54. S. J. Edmunds, N. C. Roy, D. R. Love, W. A. Laing, Kiwifruit extracts inhibit cytokine production by lipopolysaccharide-activated macrophages, and intestinal epithelial cells isolated from IL10 gene deficient mice. Cell Immunol 270, 70–79 (2011).

55. X. Chen et al., Lockd promotes myoblast proliferation and muscle regeneration via binding with DHX36 to facilitate 5′ UTR rG4 unwinding and Anp32e translation. Cell Reports 39, (2022).

56. M. Haeussler et al., Evaluation of off-target and on-target scoring algorithms and integration into the guide RNA selection tool CRISPOR. Genome Biol 17, 148 (2016).

57. K. K. H. So, et al., seRNA PAM controls skeletal muscle satellite cell proliferation and aging through trans regulation of Timp2 expression synergistically with Ddx5. 21, e13673 (2022).

58. Y. Huang et al., Large scale RNA-binding proteins/LncRNAs interaction analysis to uncover lncRNA nuclear localization mechanisms. Briefings in Bioinformatics 22, bbab195 (2021).

59. Y. Zhao et al. (Epub 2019/12/21. 10.1038/s41467-019-13598-0 PMID: 31857580).

60. Y. Li et al., Long noncoding RNA SAM promotes myoblast proliferation through stabilizing Sugt1 and facilitating kinetochore assembly. Nat Commun 11, 2725 (2020).

61. Y. Qiao et al., Nuclear m6A reader YTHDC1 promotes muscle stem cell activation/proliferation by regulating mRNA splicing and nuclear export. Elife 12, (2023).

62. C. Trapnell, D. Cacchiarelli, X. Qiu, Monocle: Cell counting, differential expression, and trajectory analysis for single-cell RNA-Seq experiments. (2018).

63. B. Langmead, S. L. Salzberg, Fast gapped-read alignment with Bowtie 2. Nature methods 9, 357–359 (2012).

64. Y. Zhao et al., Multiscale 3D genome reorganization during skeletal muscle stem cell lineage progression and aging. Sci Adv 9, eabo1360 (2023).

65. N. Servant et al., HiC-Pro: an optimized and flexible pipeline for Hi-C data processing. Genome Biol 16, 259 (2015).

66. S. S. Rao et al., A 3D map of the human genome at kilobase resolution reveals principles of chromatin looping. Cell 159, 1665–1680 (2014).

67. H. Shin et al., TopDom: an efficient and deterministic method for identifying topological domains in genomes. Nucleic acids research 44, e70–e70 (2016).

68. H. Hagege et al., Quantitative analysis of chromosome conformation capture assays (3C-qPCR). Nat Protoc 2, 1722–1733 (2007).

